# Deep skin fibroblast-mediated macrophage recruitment supports acute wound healing

**DOI:** 10.1101/2024.08.09.607357

**Authors:** Veronica M. Amuso, MaryEllen R. Haas, Paula O. Cooper, Ranojoy Chatterjee, Sana Hafiz, Shatha Salameh, Chiraag Gohel, Miguel F. Mazumder, Violet Josephson, Khatereh Khorsandi, Anelia Horvath, Ali Rahnavard, Brett A. Shook

## Abstract

Epithelial and immune cells have long been appreciated for their contribution to the early immune response after injury; however, much less is known about the role of mesenchymal cells. Using single nuclei RNA-sequencing, we defined changes in gene expression associated with inflammation at 1-day post-wounding (dpw) in mouse skin. Compared to keratinocytes and myeloid cells, we detected enriched expression of pro-inflammatory genes in fibroblasts associated with deeper layers of the skin. In particular, SCA1+ fibroblasts were enriched for numerous chemokines, including CCL2, CCL7, and IL33 compared to SCA1-fibroblasts. Genetic deletion of *Ccl2* in fibroblasts resulted in fewer wound bed macrophages and monocytes during injury-induced inflammation with reduced revascularization and re-epithelialization during the proliferation phase of healing. These findings highlight the important contribution of deep skin fibroblast-derived factors to injury-induced inflammation and the impact of immune cell dysregulation on subsequent tissue repair.

## INTRODUCTION

Skin wound healing requires intricate cell-cell communication to proceed effectively. The initial injury response generates a pro-inflammatory environment characterized by the release of inflammatory factors and the recruitment of myeloid cells (Gurtner et al., 2008, Ridiandries et al., 2018). Robust inflammation at this early stage of repair is critical, as sufficient numbers of macrophages must be recruited to become essential mediators of tissue repair (Boniakowski et al., 2018, Goren et al., 2009, Lucas et al., 2010, Mirza et al., 2009, Sawaya et al., 2020, Shook et al., 2016, Willenborg et al., 2012, Wood et al., 2014). Insufficient or excessive inflammation and macrophage recruitment prevent the wound environment from shifting to a pro-healing state that supports wound closure, precipitating the healing defects observed in conditions such as diabetes or aging (Audu et al., 2022, Eming et al., 2014, Gould et al., 2015, Joshi et al., 2020, Mirza and Koh, 2011, Mirza et al., 2014, Pang et al., 2021, Sawaya et al., 2020, Vu et al., 2022). This underscores the necessity for tight control of injury-induced inflammation and the need to define the cellular mediators of this stage of repair.

Tissue-resident cells help regulate the pro-inflammatory environment after injury. Keratinocytes are a well-established source of inflammatory cytokines and immune cell chemoattractants (Lebre et al., 2007, Villarreal-Ponce et al., 2020). Dermal adipocytes produce adipokines and lipids which can modulate wound macrophage numbers and impact wound repair (Shook et al., 2020, Zhang et al., 2019). Additionally, the adoption of a pro-inflammatory gene signature was recently established as a critical component of fibroblast activity during wound healing (Correa-Gallegos et al., 2023). These findings corroborate that injury-induced inflammation is a concerted effort from the plethora of cell types in the skin and highlight that further research is necessary to define specific roles for non-immune cells during this stage of healing.

To contextualize the changes in gene expression that occur in the different skin cell types following injury, we performed unbiased single nuclei RNA-sequencing (snRNA-seq) analysis of murine skin wounds 1-day post wounding (dpw). While keratinocytes and macrophages upregulated genes associated with immune cell chemotaxis to wounds, we found the greatest increase in pro-inflammatory gene expression within fibroblasts associated with deeper layers of the dermis. SCA1+ fibroblasts were enriched for expression of inflammatory chemokines, such as CCL2 and CCL7, and genetic ablation of *Ccl2* from fibroblasts resulted in dramatic decreases in wound macrophage and monocyte numbers during inflammation with impaired tissue healing. These findings establish fibroblasts as key mediators of injury-induced inflammation that produce essential factors for immune cell recruitment and progression through wound healing.

## RESULTS

### Stromal cells acquire a pro-inflammatory transcriptional state after injury

Diverse intercellular crosstalk pathways are transiently activated to produce pro-inflammatory, then pro-healing conditions at the injury site (Brazil et al., 2019, Eming et al., 2014, Gurtner et al., 2008, Shook et al., 2020, Shook et al., 2018, Vu et al., 2022). Single-cell RNA sequencing is a powerful identifier of cellular heterogeneity and predictive cellular function; however, cells that are fragile or tightly embedded in the extracellular matrix are frequently absent from generated datasets (Habib et al., 2017). To circumvent this limitation, we performed snRNA-seq on nuclei isolated from unwounded (UW) skin and the wound periphery from full-thickness excisional wounds (Figure 1a). Since robust populations of neutrophils, monocytes, and macrophages are present in wounds 1.5dpw (Shook et al., 2020), we isolated nuclei from tissues 24 hours (1dpw) to interrogate nuclear mRNA that could contribute to the recruitment or polarization of cells present at 1.5dpw.

**Figure 1.**
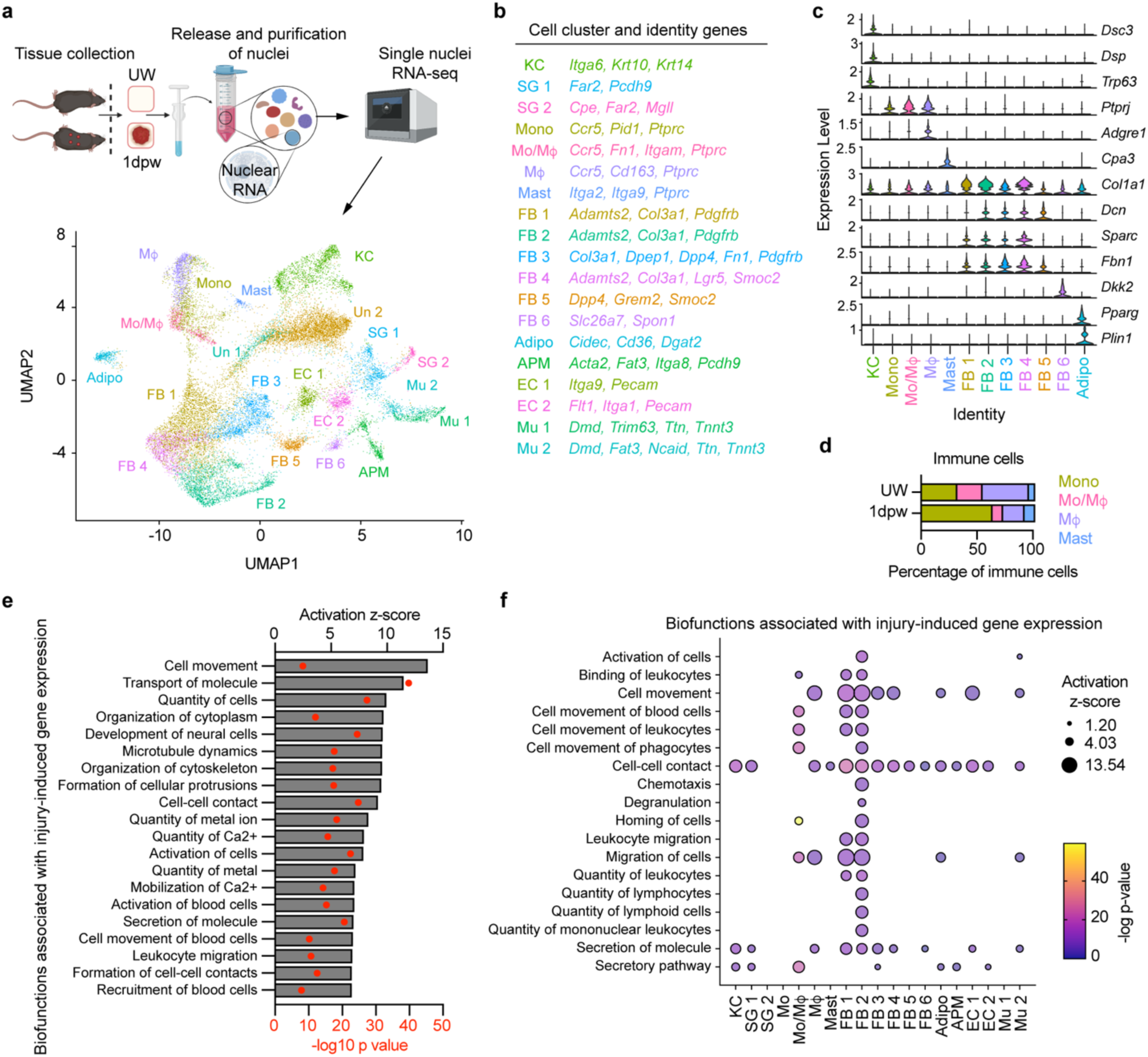
Epithelial, stromal, and immune cells adopt a pro-inflammatory gene expression profile after skin injury. (a) Schematic of nuclei harvested from uninjured and wounded skin for snRNA-seq (*n* = 2 mice per group) and Uniform Manifold Approximation and Projection (UMAP) analysis of snRNA-seq data. (b) Identification of genes for cell clusters. (c) Violin plot for genes associated with specific cellular identities. (d) Frequency of immune cell clusters in cells from unwounded (UW) skin and one-day post wound (1dpw). (e) Activated biofunctions associated with genes upregulated 1dpw. (f) Bubble matrix of Biofunctions associated with upregulated gene expression in each cell cluster 1dpw.

We identified twenty-one cell clusters from the snRNA-seq data (Figure 1a and Supplementary Figure S1a and b). Using canonical marker genes, we classified clusters as specific epithelial, immune, or stromal cell types (Figure 1a-c and Supplementary Figure S1c). Three epithelial cell clusters were identified as keratinocytes (KC) (*Dsc3, Dsp*, and *Trp63*)(Puzzi et al., 2018, Qu et al., 2018, Spindler et al., 2009) or sebaceous gland cells (SG1 and SG2) (*Far2*) (Konger et al., 2021, Rittié et al., 2016, Sundberg et al., 2018). Four immune cell populations (enriched for *Ptprc*) were classified as mast cells (*Cpa3*) (Atiakshin et al., 2022, Siddhuraj et al., 2021) and three clusters of monocytes/macrophages (*Ptprj* and *Ccr5*) (Castanheira et al., 2019, Dave et al., 2009, Oppermann, 2004, Osborne et al., 1998). Within the monocyte/macrophage groups, we identified mature macrophages (Mɸ) (high *Adgre1* and *Cd163)* (Jo et al., 2021, Pang et al., 2022), monocytes (Mono) (*Pid1)* (Lorenz et al., 2019), and a population that appeared to be immature monocyte-derived macrophages (Mo/Mɸ) (intermediate *Adgre1*). Interestingly, monocytes comprised over 50% of the immune cell population in the 1dpw samples, characteristic of monocyte influx to injury during inflammation (Figure 1d) (Crane et al., 2014). Notably, we did not detect a neutrophil cluster, though neutrophils are quickly recruited to wounds (de Oliveira et al., 2016, Shook et al., 2020). Their absence likely resulted from their generally lower RNA content and direct exposure to endogenous nucleases during tissue processing (Maraux et al., 2020, Wigerblad et al., 2022). Within the fourteen stromal cell clusters, we identified endothelial cells (EC1 and EC2) (*Pecam1*) (Tombor et al., 2021), skeletal muscle clusters (Mu1 and Mu2) (*Dmd*) (Hoffman et al., 1987), arrector pili muscle (APM) cells (*Acta2*, *Itga8*) (Ahlers et al., 2021, Fujiwara et al., 2011), adipocytes (Adipo) (*Cidec*, *Plin1*, and *Pparg*) (Kim et al., 2008, Rosen et al., 1999, Shook et al., 2020), and six fibroblast populations (FB1 - FB6) (*Col3a1*, *Dcn*, *Sparc*, and *Fbn1*) (He et al., 2023, Joost et al., 2020, Philippeos et al., 2018, Shook et al., 2020). Two clusters were not clearly enriched for markers associated with a particular cell type and were classified as unknown (Un1 and Un2).

Fibroblasts from different layers of the skin exhibit distinct gene expression profiles and perform unique functions (Correa-Gallegos et al., 2023, Driskell et al., 2013, Frech et al., 2022, Guerrero-Juarez et al., 2019, Janson et al., 2012, Jiang et al., 2020, Korosec et al., 2019, Philippeos et al., 2018). We endeavored to distinguish the six fibroblast subsets by interrogating their expression of genes associated with fibroblasts in different skin layers. One cluster (FB6) was enriched for the papillary dermis markers *Dkk2*, *Lef1*, and *Spon1* (Phan et al., 2020) (Supplementary Figure S1c). While fibroblast clusters 1-5 shared expression of the deep skin fibroblast markers *Dcn* and *Sparc* (Joost et al., 2020), FB5 had a greater enrichment for *Grem2* and *Smoc2*, suggesting they are reticular dermal fibroblasts (Phan et al., 2020) (Figure 1b and Supplementary Figure S1c). While a more granular prediction of cluster localization in the skin was not possible due to the low nuclear RNA counts, the expression pattern of these genes suggests that FB1-4 reside in the deeper layers of the dermis and the hypodermis/fascia.

Significant injury-induced changes in gene expression were defined using Tweedieverse, a differential expression tool that calculates an effect size by considering the distribution and dispersion of gene expression, thereby improving the modeling of overdispersion and zero-inflation in scRNA-seq analyses (Mallick et al., 2022). The potential biofunctions regulated by genes with a significant Tweedieverse effect size at 1dpw were determined using Ingenuity Pathway Analysis (Krämer et al., 2014). These included cell movement, leukocyte migration, secretion of molecules, and recruitment of blood cells (Figure 1e), reflecting cytokine and chemokine production that recruits myeloid cells to the wound. Indeed, predicted upstream signaling inducing these biofunctions included a variety of cytokines and their receptors (Supplementary Figure S1d), and numerous pro-inflammatory cytokines had increased expression at 1dpw (Supplementary Figure S1e).

We further investigated biofunctions associated with the 1dpw gene expression signature for each cell cluster (Figure 1f and Supplementary Figure S2a and b). Predicted biofunctions related to cell morphology, development, organization (Supplementary Figure S2a), cell signaling, molecular transport, metabolism, and cell maintenance (Supplementary Figure S2b) were shared among many populations; however, biofunctions related to immune cell recruitment were more restricted to specific cell types. Mo/Mɸ, FB1, and FB2 clusters showed the greatest predicted activation of biofunctions associated with immune cell recruitment, such as chemotaxis and quantity of leukocytes (Figure 1f).

Since keratinocytes, macrophages, fibroblasts, and adipocytes may all contribute to injury-induced inflammation (Correa-Gallegos et al., 2023, Daley et al., 2010, He et al., 2023, Lebre et al., 2007, Pang et al., 2022, Shook et al., 2020, Villarreal-Ponce et al., 2020, Zhang et al., 2019), we investigated changes in gene expression in these cells capable of influencing the immune response. Using keratinocytes and macrophages as a baseline to explore injury-induced pro-inflammatory gene expression, we observed multiple cytokines upregulated by fibroblasts after injury (Figure 2a). Most notably, *Ccl2*, *Ccl4*, *Ccl7, Il17,* and *Il33* had a significant effect size in multiple fibroblast clusters, with FB1 and FB2 having the greatest number of cytokines upregulated after injury. Fibroblasts also upregulated numerous cytokines that were not differentially expressed in keratinocytes or macrophages after injury, indicative of a prominent immune-modulatory role during early inflammation (Figure 2b). Similarly, fibroblasts upregulated many growth factors, extracellular matrix (ECM) molecules, and ECM modifiers after injury, with some shared by epithelial and immune cells (Figure 2c and Supplementary Figure S2c) and some unique to fibroblasts (Figure 2d and Supplementary Figure S2d). Interestingly, we did not detect a significant increase in cytokines or growth factors in our adipocyte population; however, low nuclear transcript counts could result in false negatives within the snRNA-seq analysis. Our gene expression analysis highlights the redundant nature of pro-inflammatory gene expression profiles among cellular populations and corroborates fibroblasts as a putative source of pro-inflammatory cues during early wound healing (Correa-Gallegos et al., 2023, Sinha et al., 2022).

**Figure 2.**
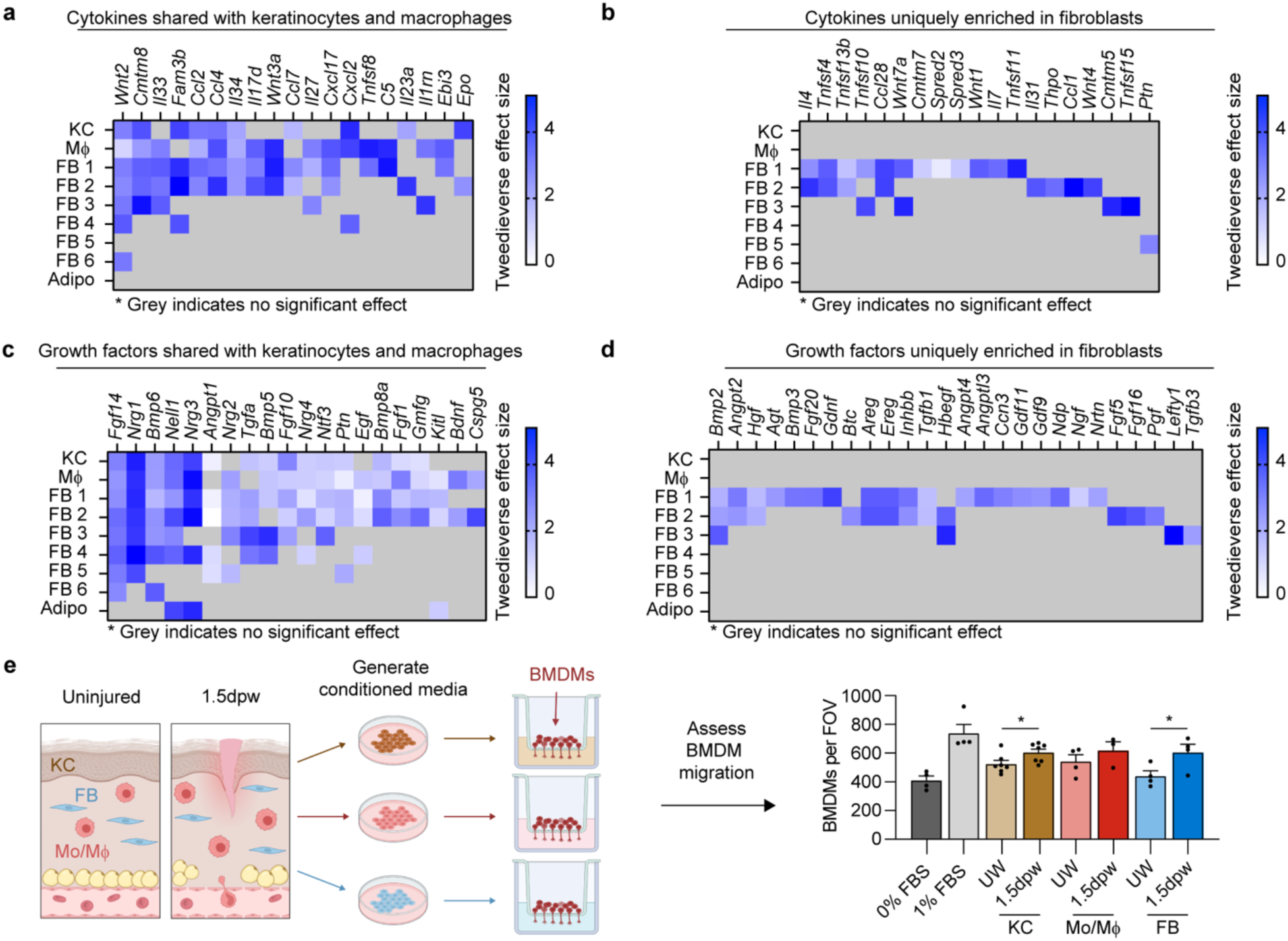
Multiple fibroblast subsets acquire a pro-inflammatory cellular state 24 hours after injury. (a-b) Cytokines with an increased effect size in multiple cell types 1dpw (a), those uniquely enriched in fibroblasts (b). (c-d) Growth factors with an increased effect size in multiple cell types 1dpw (c), those uniquely enriched in fibroblasts (d). (e) BMDM migration assay with conditioned media from primary keratinocytes, monocytes/macrophages, and fibroblasts. *n* ≥ 3 mice per group. Error bars indicate mean ± SEM. *, *p* < 0.05.

Given that many of the differentially expressed pro-inflammatory genes were chemoattractants, we evaluated the ability of keratinocytes, monocyte/macrophages, and fibroblasts to induce bone marrow-derived macrophage (BMDM) migration *in vitro*. Keratinocytes, monocytes/macrophages, and fibroblasts were isolated from uninjured skin, and 1.5dpw; 1.5dpw was chosen to allow time for nuclear RNA expression detected in the 1dpw snRNA-seq data to be converted into functional protein. Interestingly, we observed a similar injury-induced chemotactic potential of conditioned media from these cell populations (Figure 2e). These data suggest that fibroblasts can impact essential inflammatory processes, such as immune cell recruitment, to a similar level as keratinocytes and macrophages, thereby supporting their role as inflammatory mediators during healing.

### Distinct fibroblast populations have enriched expression of macrophage chemoattractants after injury

Our 1dpw snRNA-seq data indicate that fibroblasts FB1 and FB2 generate pro-inflammatory signaling molecules capable of influencing macrophage infiltration during the early stages of wound healing (Figure 1f and Figure 2a and b). To explore the spatial location of inflammatory fibroblasts in the skin, we used RNAscope to assess genes upregulated by these fibroblasts. We quantified *Ccl7* and *Il33* in *Pdgfra*+ fibroblasts at the wound periphery at 1.5dpw to capture greater quantities of mature mRNA (Figure 2a and Figure 3a). Fibroblasts significantly increased expression of both genes 1.5dpw, and *Ccl7* and *Il33*+ fibroblasts were biased towards deeper in the skin, around the panniculus carnosus and superficial fascia (Figure 3a). These results suggest that early pro-inflammatory fibroblasts reside deep in the skin and subcutaneous connective tissue, echoing previous findings that fascia fibroblasts adopt a pro-inflammatory profile after injury (Correa-Gallegos et al., 2023). Fibroblasts in the deep dermis and underlying fascia are enriched for expression of SCA1 (*Ly6a*) (Driskell et al., 2013, Lichtenberger et al., 2016), and the 5dpw transcriptomic profile of SCA1+ fibroblasts is enriched for secreted factors that can modulate macrophage migration and function (Shook et al., 2018). Several of these cytokines were identified in the 1dpw fibroblast clusters 1 and 2, including *Ccl2*, *Ccl7*, and *Il33* (Figure 2a), supporting the idea that functional diversity may exist among skin fibroblasts during inflammation.

**Figure 3.**
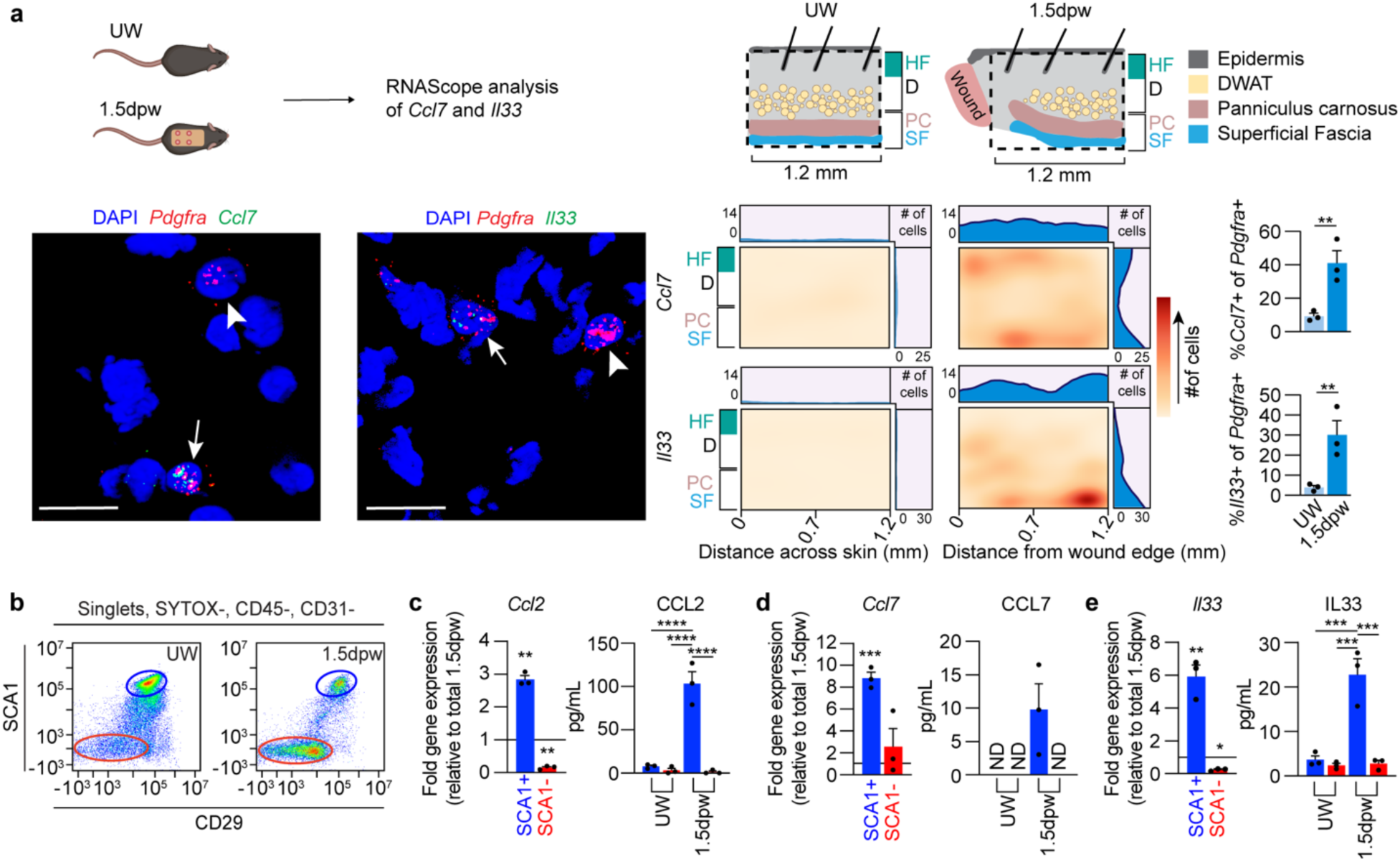
Deep skin SCA1+ fibroblasts are enriched for inflammatory gene expression. (a) RNAscope assessment of fibroblasts (*Pdgfra*+) expressing *Ccl7* or *Il33* in UW skin and at the wound periphery 1.5dpw. Images show *Pdgfra*+ fibroblasts expressing *Ccl7* or *Il33* (arrows) and lacking co-expression (arrowheads). Heatmaps show the distribution of *Ccl7* or *Il33*+ fibroblasts in the skin throughout the dermis (D), panniculus carnosus (PC), and superficial fascia (SF). “HF” indicates the depth of hair follicles into the dermis. Bar graphs show the proportion of fibroblasts positive for *Ccl7* and *Il33*. Scale bar, 15μm. *n* = 3 wound edges or UW tissue sections, each from a different mouse. (b) Flow cytometry plots of SCA1 and CD29 levels in live mesenchymal cells. (c-e) Quantitative RT-PCR relative fold change gene expression and ELISA quantification of supernatant from SCA1+ and SCA1-fibroblasts for *Ccl2*/CCL2 (c), *Ccl7*/CCL7 (d) and *Il33*/IL33 (e). Data are normalized to the expression levels in total tissue 1.5dpw. *n* = 3 mice per group. Error bars indicate mean ± SEM. *, *p* < 0.05; **, *p* < 0.01; ***, *p* < 0.001; ****, *p* < 0.0001.

To further validate that deeper fibroblasts are enriched for pro-inflammatory gene expression after injury, we isolated lineage-negative stromal cells (CD45-, CD31-) based on SCA1 and CD29 surface levels at 1.5dpw (Figure 3b and Supplemenatary Figure S3a). SCA1+ fibroblasts were significantly enriched for expression of *Ccl2*, *Ccl7*, *Cxcl1, Cxcl12, Il6,* and *Il33* at 1.5dpw compared to the expression level in the entire wound bed (Figure 3c-e and Supplementary Figure S3b). The expression level of these genes in the SCA1-population was not significantly different or was significantly lower (Figure 3c-e and Supplementary Figure S3b). Secreted protein analysis corroborated the gene expression data, indicating that SCA1+ fibroblasts generate significantly more CCL2, CCL7, and IL33 than SCA1-cells, as well as significantly more than SCA1+ fibroblasts from uninjured skin (Figure 3c-e). Overall, these results support that stromal cells, particularly SCA1+ fibroblasts, are a key local source of inflammatory factors during early injury-induced inflammation.

### Fibroblast-derived CCL2 is required for injury-induced macrophage recruitment

Signaling through CCR2 is essential for monocyte and macrophage trafficking to the wound (Boniakowski et al., 2018, Willenborg et al., 2012, Wood et al., 2014). The upregulation of fibroblast *Ccl2* in our snRNA-seq data and the enriched expression and secretion of *Ccl2*/CCL2 in SCA1+ fibroblasts implicate CCL2 as a key mediator of fibroblast-macrophage communication during injury-induced inflammation. To assess how fibroblast-derived CCL2 impacts immune cell numbers in the wound, we crossed *Pdgfra*CreER mice (Chung et al., 2018) with *Ccl2* floxed mice (Shi et al., 2011) to genetically target *Ccl2* in fibroblasts. SCA1+ fibroblasts isolated from *Pdgfra*CreER+; *Ccl2*^fl/fl^ (fibroblast conditional knock-out, FBcKO) mice at 16 hours post-wounding (hpw) and 1.5dpw displayed reduced *Ccl2* expression compared to *Pdgfra*CreER-; *Ccl2*^fl/fl^ (control) mice (Supplementary Figure S3c and d).

We evaluated the effect of fibroblast-derived CCL2 on immune cell numbers during the inflammatory phase at 1.5dpw and 3dpw (Figure 4a and b). At 1.5dpw, FBcKO mice displayed greater than a 50% reduction in wound macrophage and monocyte numbers compared to control mice (Figure 4c), suggesting a profound defect in myeloid cell trafficking to the wound. Neutrophils and total CD45+ immune cells were also slightly reduced. Immunostaining for the macrophage marker F4/80 in 1.5dpw tissue sections revealed that the reduction in macrophages was limited to the wound bed in FBcKO mice (Figure 4d-g). SCA1+ fibroblasts from FBcKO mouse wounds possessed diminished *in vitro* macrophage chemotactic potential compared to fibroblasts from control mice (Supplementary Figure S3e). Interestingly, by 3dpw, the trends in immune cell numbers had switched, as FBcKO mice tended to have higher numbers of immune cells, macrophages, and monocytes per wound compared to control mice, though the data did not reach statistical significance (Figure 4h). To explore this phenomenon, we evaluated gene expression in SCA1+ and SCA1-fibroblasts for the chemokines *Ccl2* and *Ccl7*, which also signals through CCR2 (Tsou et al., 2007, Willenborg et al., 2012), at 16hpw and 1.5dpw (Supplementary Figure S3c and d). At 16hpw, *Ccl2* expression was lower in the total wound and SCA1+ fibroblasts from FBcKO mice compared to controls, and *Ccl7* expression was not significantly different between the groups. However, by 1.5dpw, total wound *Ccl2* expression was comparable, and *Ccl7* expression was significantly higher in the SCA1+ fibroblasts from FBcKO mice. Surprisingly, we did not observe any change in *Ccl2* or *Ccl7* expression in SCA1-fibroblasts in either mouse strain. Together, these results suggest that loss of CCL2 production from the SCA1+ fibroblasts impacts wound myeloid cell numbers in the earliest stages of inflammation, but compensatory increases in chemokine expression eventually allow for robust recruitment of myeloid cells.

**Figure 4.**
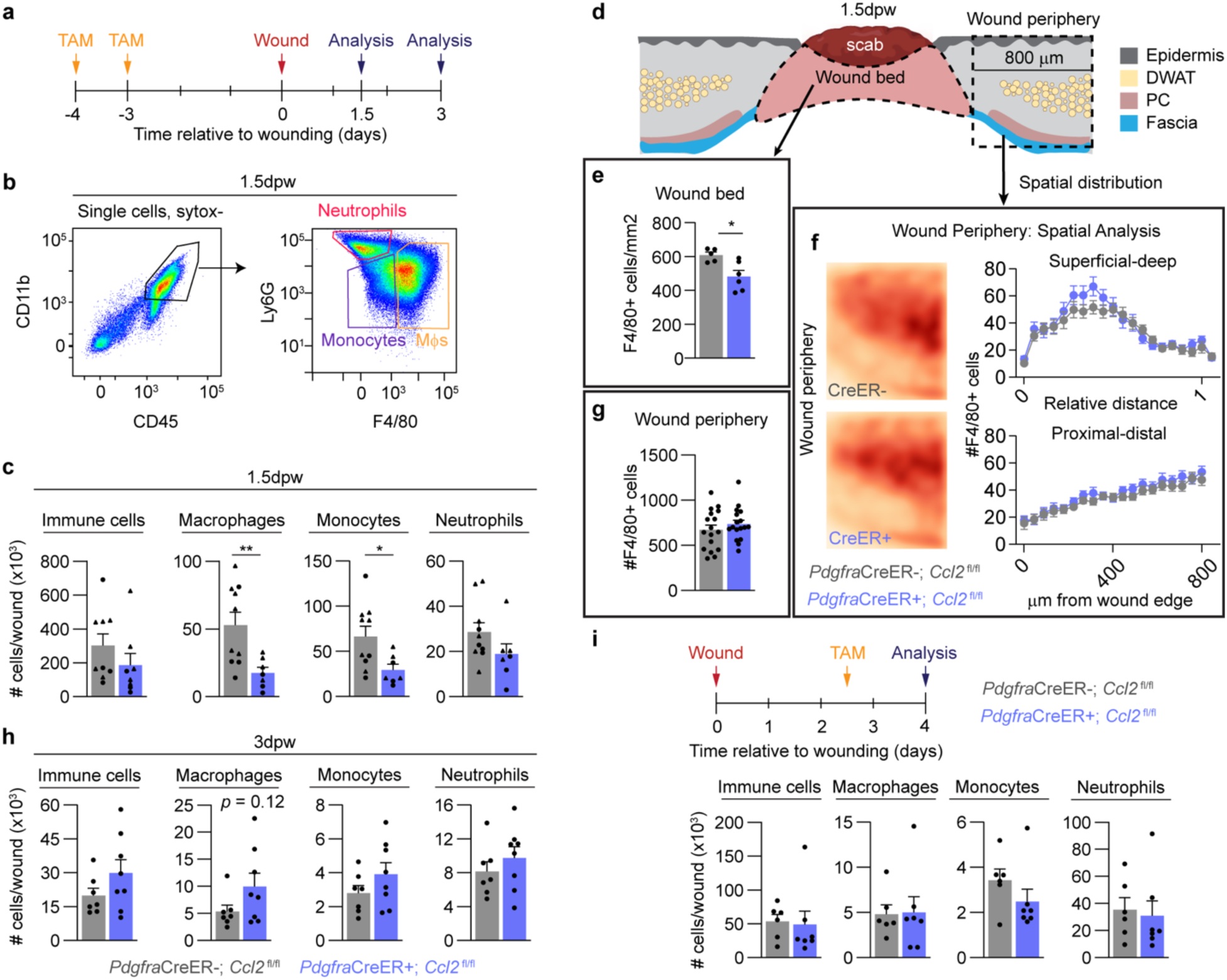
Fibroblast-derived CCL2 supports macrophage numbers during the inflammation phase of wound healing. (a) Paradigm to induce *Ccl2* deletion in *Pdgfra*-expressing cells and analysis time points. (b) Gating strategy for immune cell populations in wound beds. (c) Flow cytometry quantification of immune cell subsets in FBcKO and control mice. *n* ≥ 7 mice per group. (d) Schematic of spatial tissue analysis of macrophages (F4/80+) in FBcKO and control mice. (e) Quantification of F4/80+ cells in the wound centers. *n* ≥ 5 wounds from 4 different mice per group. (f-g) Quantification of F4/80+ cells at the wound periphery. (f) Heatmaps and graphed distribution of F4/80+ cells along the superficial-deep and proximal-distal axes. (g) Total numbers of F4/80+ cells in the wound periphery. *n* ≥ 16 wound edges from 4 mice per group. (h) Flow cytometry quantification of immune cell subsets at 3dpw. *n* = 7 mice per group. (i) Timeline and immunophenotyping quantification for tamoxifen administration to FBcKO and control mice at the end of the inflammatory phase. *n* ≥ 6 mice per group. Triangles and circles delineate female and male mice, respectively. Error bars indicate mean ± SEM. *, *p* < 0.05; **, *p* < 0.01.

We also explored whether fibroblast-derived CCL2 supports the continued recruitment of monocytes and macrophages to the wounds after peak inflammation. Knockout of *Ccl2* in fibroblasts at 2.5dpw did not alter the number of immune cells present in wounds 4dpw (Figure 4i). This further emphasizes the role of SCA1+ fibroblast-derived CCL2 early after injury and suggests that fibroblasts play an essential role in the rapid, early recruitment of innate immune cells to the wound.

### Skin wound healing is impaired in fibroblast *Ccl2* knock-out mice

Directly reducing macrophage numbers during the inflammation phase delays revascularization, re-epithelialization, and granulation tissue formation (Goren et al., 2009, Lucas et al., 2010, Mirza et al., 2009, Shook et al., 2016, Willenborg et al., 2012). To determine if the reduction in wound bed macrophages in FBcKO mice affects downstream wound healing, we examined multiple reparative processes in wounds 5dpw. Immunostaining tissue sections from the center of wound beds with ITGA6 (Yan et al., 2022) revealed that re-epithelialization of wound beds was reduced in FBcKO mice, with fewer wounds becoming fully re-epithelialized or closed at 5dpw compared to controls (Figure 5a). Wound width was also slightly higher in the FBcKO group, suggesting that these mice may have altered wound contraction (Chen et al., 2015, Lucas et al., 2010, Parfejevs et al., 2018).

**Figure 5.**
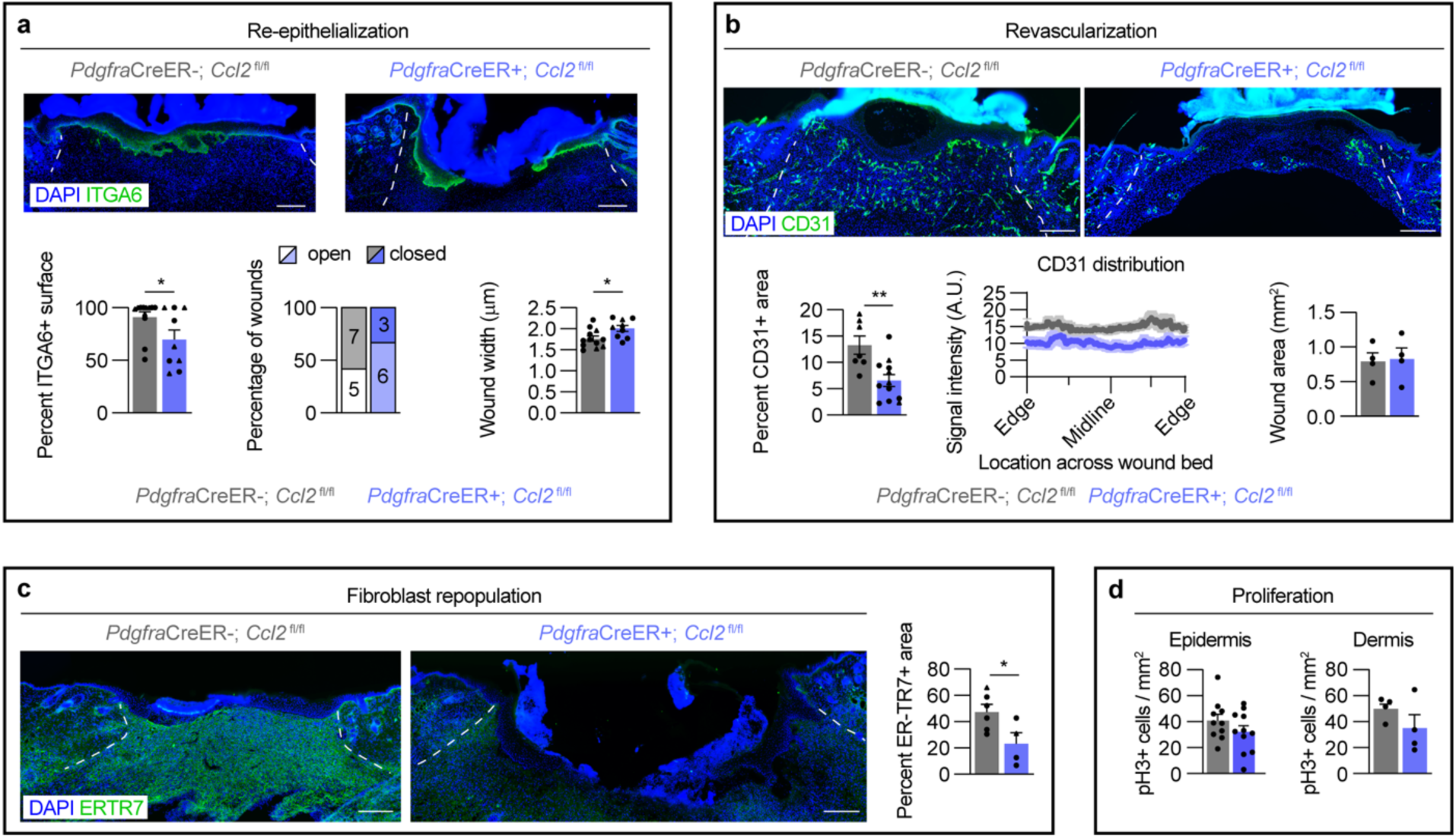
Epidermal and dermal repair are delayed in fibroblast *Ccl2* knock-out mice. (a) Images of wound beds of FBcKO and control mice 5dpw immunostained for ITGA6 and DAPI. Graphs show the percentage of ITGA6+ surface (left), the frequency of open or closed wounds (midline), and the wound width (right). n ≥ 9 wounds from n ≥ 6 mice per group. (b) Images of wound beds of FBcKO and control mice 5dpw immunostained for CD31 and DAPI. Graphs show the CD31+ area (left), the average distribution of CD31 signal intensity from the wound edge to the center (midline), and the total wound bed area (right). n ≥ 4 wounds from n ≥ 3 mice per group. (c) Images and quantification from wound beds of FBcKO and control mice 5dpw immunostained for ER-TR7 and DAPI. n ≥ 4 wounds from n ≥ 3 mice per group. (d) Quantification of pH3+ cells in the epidermal (left) and dermal (right) compartments of wound beds 5dpw in FBcKO and control mice. n ≥ 4 wounds from n ≥ 3 mice per group. Scale bars are 250µm, and white lines delineate wound edges. Data points indicate individual wounds. Triangles and circles delineate female and male mice, respectively. Error bars indicate mean ± SEM. \**p* < 0.05, \*\**p* < 0.01. A.U. Arbitrary units of fluorescence.

We further investigated if FBcKO mice displayed delayed healing in the dermal compartment. Compared to control mice, revascularization and fibroblast repopulation were reduced by 50% throughout the wound bed, though the average wound bed area was the same (Figure 5b and c). Notably, we did not observe a difference in the number of phospho-histone H3+ (pH3+) mitotic cells in the epidermis or dermis 5dpw (Figure 5d). Overall, the significant decrease in wound myeloid cell numbers during inflammation and subsequent healing delay observed in the fibroblast *Ccl2* knockout mice highlights that fibroblast pro-inflammatory signaling is essential for timely progression through the healing process.

## DISCUSSION

Mesenchymal cells are increasingly recognized as key coordinators of immune cell activity at homeostasis and after injury (Krausgruber et al., 2020, Zhou et al., 2022). Fibroblasts in the heart and lung activate inflammatory cytokine expression and recruit immune cells in acute injury or infection (Boyd et al., 2020, Choi et al., 2004, Kawaguchi et al., 2011, Mouton et al., 2019, Yu et al., 2013). Dermal fibroblasts activate cytokine expression in response to toll-like receptor stimulation or treatment with inflammatory factors (Al-Rikabi et al., 2021, Kitanaka et al., 2019, Ploeger et al., 2013, Yao et al., 2015). While relatively few studies investigate dermal fibroblasts *in vivo* during the early stages of wound healing, inflammatory fibroblasts have been observed at 1dpw (Hu et al., 2023), and an inflammatory phenotype was recently described as an essential step in the development of wound myofibroblasts (Correa-Gallegos et al., 2023). Here, we confirm that fibroblasts are potent modulators of inflammation early after skin injury, upregulating a myriad of factors with functional importance in wound healing (Figure 6). This finding was emphasized by the fact that fibroblast-induced macrophage migration was comparable to keratinocytes and monocytes/macrophages, two well-studied cellular sources of chemotactic factors (Guenin-Mace et al., 2023, Pang et al., 2022, Ridiandries et al., 2018, Roupé et al., 2010, Villarreal-Ponce et al., 2020). While we focused predominantly on the role of fibroblasts in macrophage/monocyte recruitment, we also observed upregulation of factors involved in neutrophil chemotaxis (*Cxcl2*) (De Filippo et al., 2013, Sawant et al., 2021) and macrophage polarization (*Il4, Il33*, and *Il34*) (Chen et al., 2023, He et al., 2017, Oh et al., 2023, Zhuang et al., 2023) after injury. Given that fibroblasts may regulate neutrophil migration and macrophage polarization in other inflammatory contexts (Ferrer et al., 2017, Le Fournis et al., 2021, Williams et al., 2021, Witowski et al., 2009), these findings suggest that fibroblasts perform multiple conserved roles in innate immune cell inflammation across tissues.

**Figure 6.**
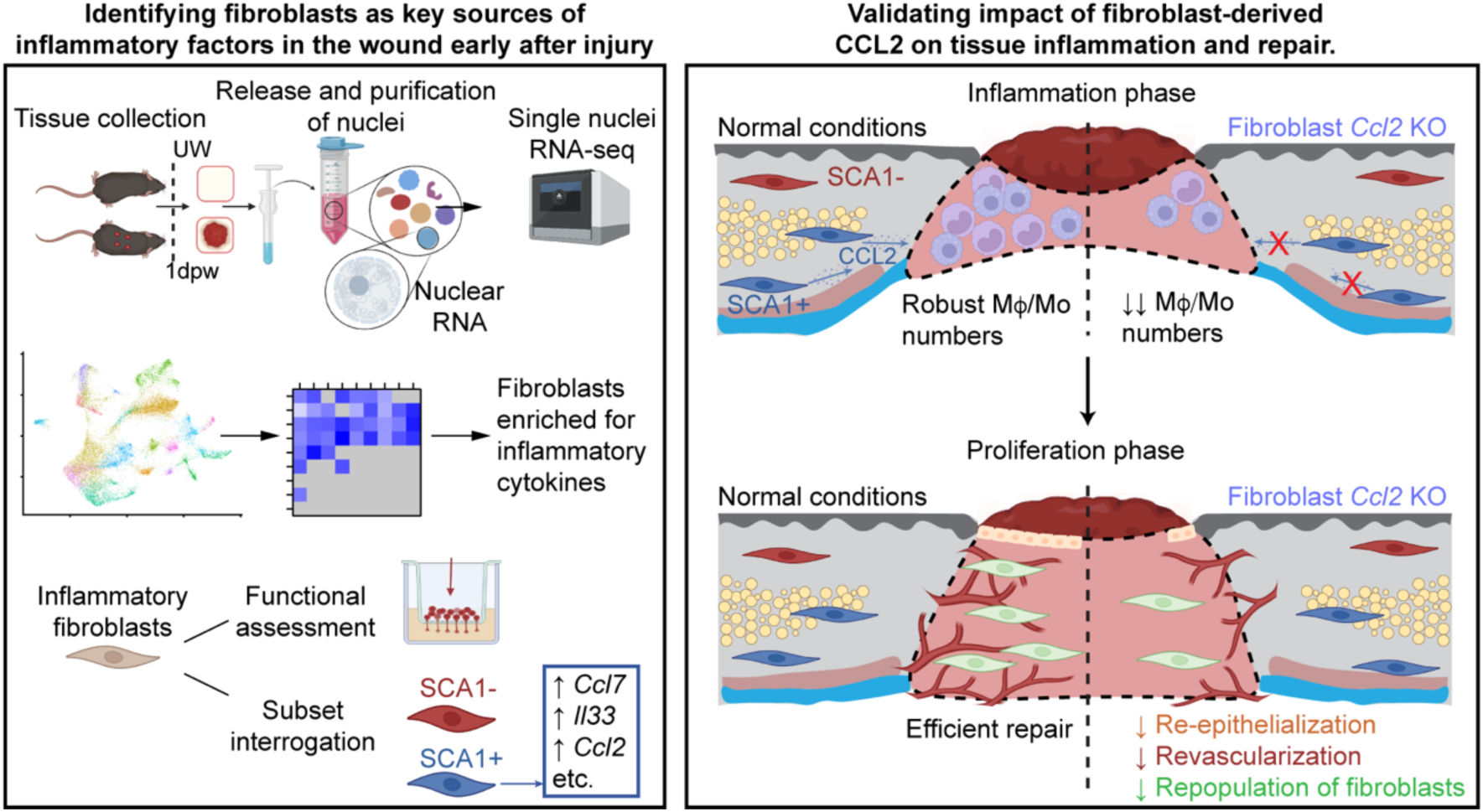
Fibroblasts in the deep skin produce inflammatory factors during the early injury response that are critical for immune cell recruitment and subsequent tissue repair.

In the skin, robust myeloid cell infiltration, particularly of macrophages, is essential for successful repair, as evidenced by the delayed and impaired healing observed when early immune cell infiltration is impeded (Boniakowski et al., 2018, Goren et al., 2009, Lucas et al., 2010, Mirza et al., 2009, Sawaya et al., 2020, Shook et al., 2016, Willenborg et al., 2012, Wood et al., 2014). While many chemokines participate in generating a chemotactic gradient to draw immune cells to the wound, CCL2 has been consistently identified as a key factor for monocytes/macrophages in this process (Willenborg et al., 2012, Wood et al., 2014). *Ccl2* expression was previously observed in dermal fibroblasts after injury (Correa-Gallegos et al., 2023, Shook et al., 2018). Here, we functionally validated that fibroblast-derived CCL2 is essential for robust wound monocyte/macrophage numbers and timely wound closure and repopulation. Given that CCL2 also drives monocyte/macrophage recruitment after injury in other tissues such as the heart, lungs, and skeletal muscle (Dewald et al., 2005, Lu et al., 2011, Pappritz et al., 2018, Suresh et al., 2012), it will be interesting to see whether fibroblasts are also an important source of this chemokine in other injury settings and expand our understanding of critical fibroblast-macrophage communication across tissues and conditions (Zhou et al., 2022).

Skin fibroblasts are highly heterogeneous, with differences in genetic lineage or spatial location in the skin often underscoring unique functions of different subsets (Abbasi et al., 2020, Correa-Gallegos et al., 2023, Driskell et al., 2013, Foster et al., 2021, Guerrero-Juarez et al., 2019, Korosec et al., 2019, Leavitt et al., 2020, Phan et al., 2020, Rinkevich et al., 2015). We observed upregulation of pro-inflammatory factors in fibroblasts from deep in the skin, particularly those expressing SCA1 (Driskell et al., 2013, Lichtenberger et al., 2016, Rognoni et al., 2016), but not the regeneration-associated papillary fibroblasts (Frech et al., 2022, Phan et al., 2020). This echoes findings that fascia-resident fibroblasts exhibit a pro-inflammatory profile after inflammation (Correa-Gallegos et al., 2023). SCA1+ fibroblasts were previously predicted to influence inflammation and chemotaxis based on their gene expression in uninjured skin (Philippeos et al., 2018). These fibroblasts share markers of human fibroblasts found in the deep dermis, which respond robustly to inflammatory cues after isolation from healthy skin (Korosec et al., 2019, Philippeos et al., 2018). Moreover, SCA1 also delineates two major fibroblast populations in the heart, and SCA1+ cardiac fibroblasts are enriched for expression of *Ccl2* and other chemokines compared to SCA1-fibroblasts after heart injury (Chen et al., 2018, Farbehi et al., 2019). This implies that subset-specific inflammatory fibroblast signaling may be conserved across tissues. Why these subsets are enriched for this function requires further investigation.

Dysregulated inflammatory signaling from fibroblasts has been associated with various pathologies in the skin and other organs. In diabetic and aged skin wounds, fibroblasts express decreased levels of immune cell chemoattractants, yet maintain excessive expression of other factors that drive chronic inflammation and poor healing (Al-Rikabi et al., 2021, Januszyk et al., 2020, Mahmoudi et al., 2019, Vu et al., 2022, Wall et al., 2008). Similarly, sustained inflammatory signaling and even CCL2 production from fibroblasts have been associated with fibrosis and scar formation in the skin and other organs (Bhattacharyya et al., 2018, Burke et al., 2010, Chen et al., 2018, Correa-Gallegos et al., 2023, Foster et al., 2021, Jiang et al., 2020, Paish et al., 2018, Sinha et al., 2022, Wong et al., 2011, Wu et al., 2014, Zhang et al., 2012). Our findings suggest that a “healthy” level of fibroblast inflammatory gene expression exists which is critical for appropriate immune responses early after injury. Identifying factors that initiate, suppress, or prolong this inflammatory state in fibroblasts will be fundamental to addressing healing pathologies associated with insufficient or excessive inflammation.

## MATERIALS AND METHODS

### Animals

Animal maintenance and experiments were conducted under the guidance of George Washington University’s Institutional Animal Care and Use Committee (IACUC). C57/Bl6 mice were purchased from Jackson Laboratories (JAX, Bar Harbor, ME, USA, strain 000664). *Pdgfra*CreER; *Ccl2*^fl/fl^ mice were generated by crossing B6.129S-*Pdgfra*tm1.1(cre/ERT2)Blh/J (Jackson Laboratories, Bar Harbor, ME, USA, strain 032770) with B6.Cg-*Ccl2^tm1.1Pame^*/J (Jackson Laboratories, Bar Harbor, ME, USA, strain 016849). Tamoxifen was administered to *Pdgfra*CreER; *Ccl2* ^fl/fl^ mice topically to shaved back skin (2x applications of 100 µL of 5 mg/mL in ethanol, Sigma-Aldrich T5648) for all experiments except Figure 3d-f, Figure 3i, and Supplementary Figure S3c and d. For these experiments, tamoxifen was administered by intraperitoneal (IP) injection (2× 100 µL injections of 30 mg/mL in sesame oil, Sigma-Aldrich S3547-250ML). All experimental procedures were approved in accordance with George Washington University’s Institutional Animal Care and Use Committee (IACUC). Male mice were used for all experiments unless otherwise noted in figure legends.

### Dorsal skin excision

Male and female mice in the telogen phase of the hair cycle were used (7-10 weeks old). For wounding experiments, mice were anesthetized with isoflurane and inflicted with two or four, excisional wounds 4-6 mm apart on shaved back skin by a 4 mm biopsy punch (Miltex).

### Isolation of single nuclei for RNA-sequencing

Methods were adapted from Li, et al 2022 (Li and Humphreys, 2022) and Habib, et al 2017 (Habib et al., 2017). To make nuclei lysis buffer 0 (NLB0), one cOmplete ULTRA tablet (Roche 05892791001) was dissolved per 10mL Nuclei EZ Lysis Buffer (Sigma-Aldrich Nuc-101). To make Nuclei lysis buffer 1 (NLB1), 10μL RNasin Plus ribonuclease inhibitor (Promega, N2615) and 10μL SUPERaseIN RNase Inhibitor (Invitrogen AM2694) were added per 4 mL NLB0. To make NLB2, 4μL RNasin Plus ribonuclease inhibitor and 4μL SUPERaseIN Rnase Inhibitor were added per 4mL NLB0. All steps were performed on ice. Briefly, skin was collected from male non-wounded and 24 hours post-wounding C57BL/6 mice and minced to approximately 1mm^2^ in NLB1. Minced tissue was transferred to KONTES dounce tissue grinders (Kimble Chase KT885300-0002) with an additional 1mL of NLB1. The loose pestle (A) was applied 22 times before passage through 200 µm pluriStrainer (pluriSelect 43-50200). The tight pestle (B) was applied 12 times, then the homogenate was incubated for 5 minutes with an additional 2mL NLB1. Then the isolated nuclei were passed through a 40 µm pluriStrainer (pluriSelect 43-50040), centrifuged at 500 x g, 5 min, 4°C and resuspended in 4mL NLB2. Resuspended samples were incubated on ice for 5 minutes, then centrifuged. Samples were resuspended in suspension buffer (NSB; 1µL RNasin Plus/mL 2% BSA in DPBS). Nuclei were washed by centrifugation and resuspension 4 times before a final resuspension in 250µL NSB and filtration through a 5µm strainer. 50µL were taken for assessment and counting of nuclei, and the remainder were diluted according to 10X protocol targeting 10,000 nuclei per sample.

### Construction of 10X Genomic Single Cell 3’ RNA-seq libraries and sequencing

Immediately after isolation, 2 wounded and 2 uninjured samples were loaded into 10X genomics Next GEM G chips (10X Genomics 2000177) with Next GEM Single Cell 3’ Gel Beads v3.1 (10X Genomics 2000164) for subsequent lysis, cDNA synthesis, and amplification. Library prep was performed using Single Index Kit T Set A (10X Genomics 2000240). Samples were balanced and sequenced by Genewiz (now Azenta), with the aim of 50,000 reads per cell. All 4 samples were pooled in one lane on an Illumina NovaSeq 6000 instrument, which averages 2 billion reads per lane. Data are available at GEO: GSE265996 (reviewer token: ovwriiaqdlkztur).

### Quantification and integration of snRNA-seq data

Data preprocessing and analysis were performed using Cell Ranger 5.0.1 (10x Genomics) (Zheng et al., 2017). Briefly, Cell Ranger’s count utility was used to align the raw sequencing reads to mm10 and to generate feature-barcode matrices for downstream analyses. To accommodate the characteristics of nuclear RNA, which includes a significant proportion of unspliced pre-mRNA, the “--include-introns” flag was used. Quality metrics, such as the total number of detected genes per nucleus, the proportion of reads mapping to the nuclear genome, and the distribution of reads across intronic and exonic regions were assessed to filter out low-quality nuclei.

Downstream quality control and data integration were performed in R using Seurat (Satija et al., 2015, Stuart et al., 2019). Samples were filtered based on the number of features, RNA count, and percentage of mitochondrial genes per nucleus, following established quality control methods (Luecken and Theis, 2019, Osorio and Cai, 2021). Due to variations in sample quality, filtering parameters were determined for each sample (Luecken and Theis, 2019). For Unwounded Samples #1 and #2 the nFeature_RNA > 200, and nFeature_RNA < 2500 with nCount_RNA < 8000, and the percent.mt < 1. For 1dpw Sample #1 the nFeature_RNA > 200, and nFeature_RNA < 3000 with nCount_RNA < 6000 and the percent.mt < 1.5. For 1dpw Sample #2, the nFeature_RNA > 200, and nFeature_RNA < 1750 with nCount_RNA < 2250, and the percent.mt < 1. Data normalization was performed using sctransform v0.4.1 (Hafemeister and Satija, 2019). Following normalization, integration was performed using Seurat’s FindIntegrationAnchors() and IntegrateData() functions using the default setting of 30 canonical correlation analysis dimensions (Hao et al., 2021). Twenty-one cell clusters were subsequently identified with Seurat’s standard RunPCA(), FindClusters(), and FindNeighbors() functions, using the first 30 principle components and a resolution of 0.8 (Xu and Su, 2015). Seurat’s FindAllMarkers() function was used to identify genes associated with each cluster.

### Gene Expression and Pathway Enrichment Analysis

To measure differences in gene expression between the unwounded and injured conditions in the snRNA-seq dataset, we employed Tweedieverse v0.0.1 (Mallick et al., 2022), targeting a false discovery rate of 0.05 and using the Benjamini-Hochberg procedure to report significance. Significant changes in gene expression are reported as the Tweedieverse effect size. Pathway enrichment analysis to predict Biofunctions associated with changes in gene expression after injury was performed using Ingenuity Pathway Analysis (Krämer et al., 2014).

### Flow cytometry and sorting

Following euthanasia with CO_2_ and cervical dislocation, mouse back skins or wound beds were excised and minced. Samples were digested in RPMI (Gibco 61870-036) with 0.25mg/mL Liberase TM (Roche 05401127001), 1mM Sodium Pyruvate (Gibco 11360070), 1% Antibiotic-Antimycotic (anti-anti; Gibco 15240062), 50U/mL Deoxyribonuclease I (DNase I; Worthington LS002006), 2% MEM Non-Essential Amino Acids (Gibco 11140050), and 25mM HEPES buffer (Gibco 15630080). Samples were incubated in a shaker incubator at 37°C for 90 minutes to release single cells. Tissue samples were consecutively filtered through 70 µm and 40 µm strainers. Single-cell suspensions from tissue digestion or epidermal isolation were resuspended in FACS staining buffer (1% BSA in phosphate-buffered saline (PBS; Gibco 14190-250) with 2nM EDTA and 25U/mL DNase I) and immunostained for 20 minutes on ice.

To assess mesenchymal heterogeneity, cells were stained with the following antibodies: CD45-APC-Cy7 (Biolegend, 1:1000), CD31-APC-Fire-750 (Biolegend, 1:500), CD29-Alexa700 (Biolegend, 1:500), EpCAM-APC-Fire-750 (Biolegend, 1:500), and SCA1 (LY6A/E)-BV650 (Biolegend, 1:1500). For myeloid cell immunophenotyping, cells were stained with the following antibodies: CD45-APC-Cy7 (Biolegend, 1:1000), CD11b-Alexa700 (Biolegend, 1:500), F4/80-eFluor450 (eBioscience, 1:200), and Ly6G-BV785 (Biolegend, 1:200) or -PE-Cy7 (Biolegend, 1:500). Wound monocytes were defined as CD45+; CD11b+; F4/80-; Ly6G-cells. Macrophages were defined as CD45+; CD11b+; F4/80+; Ly6G-cells. Neutrophils were defined as CD45+; CD11b+; F4/80-; Ly6G+ cells. For experiments where fibroblasts and the monocyte/macrophage population were isolated from the same wounds, cells were stained with CD45-FITC (Biolegend, 1:1000), CD31-APC-Fire-750 (Biolegend, 1:500), CD11b-Alexa700 (Biolegend, 1:500), EpCAM-APC-Fire-750 (Biolegend, 1:500), Ly6G-PE-Cy7 (Biolegend, 1:500), and SCA1-BV650 (Biolegend, 1:1500). Wound monocyte/macrophages were defined as CD45+CD11b+Ly6G-cells.

For keratinocyte isolation, total back skin or 1-2mm of skin surrounding wounds was harvested from the backs of the mice. Fat was scraped from the bottom of the skin using a blunt scalpel. The skin samples were placed dermis side down in a sterile culture dish and rinsed in PBS without Ca2+ or Mg2+ (Gibco 14190-250). Samples were suspended in 0.25% Trypsin-EDTA (Gibco 25200072) and incubated for 75 minutes at 37°C. Following incubation, hair was scraped from the skin using curved forceps and the scalpel. Samples were broken down with the scalpel and triturated with a 25mL pipet for ∼50 seconds. Samples were filtered through 70μm and 40μm strainers and cells were stained with CD45-APC-Cy7 (Biolegend, 1:1000) and EpCAM-APC (Biolegend, 1:100) to isolate CD45-; EpCAM+ cells.

To exclude dead cells, samples were incubated with Zombie Aqua (Biolegend 423102, 1:1000) at room temperature for 20 minutes before antibody staining or Sytox Blue or Sytox Green (Invitrogen S34857 or S34860, 1:1000) was added before analysis. Analysis was performed on a Celesta (BD Biosciences) or CytoFLEX S (Beckman Coulter Life Sciences), and sorting was performed on a BD Influx (BD Biosciences) or a Sony SH800 (SONY). Cells were sorted into 10% fetal bovine serum (FBS; Gibco A4766801), 1% anti-anti (Gibco 15240062) in DMEM (ATCC 30-2002), and flow cytometry analysis was performed using FlowJo Software (FlowJo).

Additional information on antibodies used for flow cytometry can be found in Supplementary Table S1.

### Quantitative Real-Time PCR

FACS-purified cells from uninjured skin or wound beds were lysed in TRIzol (Invitrogen 15596018); RNA was isolated from the aqueous phase and purified using the RNeasy Micro kit (Qiagen 74004). The Superscript IV kit (Invitrogen 18-090-200) was used to generate cDNA proportionally to the isolated RNA. Quantitative real-time PCR was performed using PowerTrack SYBR Green master mix (Applied Biosystems A46111) run on a BioRad CFX384 with a C1000 thermocycler. Gene-specific primer sequences are listed in Supplementary Table S2. All reported values reflect the levels of target mRNA normalized to β-actin. mRNA levels are the average of biological samples (*n* ≥ 3) calculated from technical qPCR triplicates.

### Enzyme-linked immunosorbent assay (ELISA)

Skin wounds were digested and stained for flow cytometry as described above, and cell populations were sorted on a BD Influx. Cells were plated in a 24-well tissue culture plate (Corning COSTAR 3526) at a density of at least 36,000 cells/mL in DMEM with 10% FBS and 1% anti-anti and incubated at 37°C for 2.5 hours. Cell supernatant was harvested after incubation. ELISA kits were used to measure the concentration of CCL2 (R&D Systems MJE00B), CCL7 (Invitrogen BMS6006INST), and IL33 (R&D Systems M3300). Briefly, supernatant (two technical replicates per sample) was incubated on the provided plate, along with a set of standard dilutions. After washing the wells, conjugate specific to the analyte was added, followed by further incubation, washing, and addition of substrate. The optical density of each well was determined using a BioTek Synergy H1 Hybrid Reader set to 450nm. Cell supernatant was either diluted prior to the ELISA or the detection values were normalized to the lowest seeding density if the number of fibroblasts seeded differed between biological replicates.

### Bone Marrow Isolation and Macrophage Differentiation

Mice were euthanized with CO2 and cervical dislocation. The femur and tibia were isolated from both hind limbs. After cleaning, the bones were cut at both ends, and bone marrow was flushed out with sterile RPMI and a 25-gauge needle. Red blood cells were lysed with 2mL of ACK lysis buffer (150mM NH_4_Cl, 10mM KHCO_3_, 0.1mM Na_2_EDTA). After buffer neutralization with RPMI plus 5% FBS, cells were passed through a 70μm filter. Cells were plated at a density of 3×10^5^ cells/mL in 10mL of RPMI plus 30% L929 conditioned media, 10% FBS, and 1% anti-anti in a 10cm petri dish. On day 3 or day 4 after plating, an additional 10mL of this media was added to the plates. BMDMs were used for experiments on day 6 or day 7. L929 cells (MilliporeSigma 85011425-1VL) were cultured and conditioned media was harvested as previously described (Weischenfeldt and Porse, 2008).

### Transwell migration assay

#### BMDM migration in response to Fibroblasts, Myeloid cells, and Keratinocytes

Fibroblasts (CD45-CD31-EpCAM-), myeloid cells (CD45+CD11b+Ly6G-), and keratinocytes (CD45-EpCAM+) were sorted from uninjured skin and wound samples 1.5dpw after skin digestion (fibroblasts and macrophages) or epidermal isolation (keratinocytes) as described above. Cells were plated at a density of ∼40,000 cells/mL in serum-free RPMI with 1% anti-anti immediately after sorting and incubated at 37°C with 5% CO2 overnight. Conditioned media was harvested from the cells and spun down to remove any debris before storage at −80°C until use.

Twenty-four hours prior to the migration assay, BMDMs were serum starved in RPMI. Serum-starved BMDMs were lifted from the plate with a cell scraper in cold PBS and resuspended in RPMI at 1×10^6^ cells/mL. 700uL of conditioned media from the cell types of interest was added to the bottom of a 24-well tissue culture plate. 100uL of the BMDM solution was added to a transwell insert with 8μm pore diameter (MilliporeSigma PTEP24H48) submerged in the conditioned media. Wells with RPMI with 1% anti-anti and no FBS (0% FBS) or 1% FBS (1% FBS) were used for internal negative and positive controls for the assay, respectively. Assay plates were incubated overnight at 37°C with 5% CO2.

#### Comparing myeloid cell migration between PdgfraCreER+; and PdgfraCreER-; Ccl2^fl/fl^ mice

SCA1+ fibroblasts and monocytes/macrophages were sorted from the same 1.5d wounds. SCA1+ fibroblasts were plated at a density of 36,000 cells/mL in the bottom of a 24-well tissue culture transwell insert plate (Corning COSTAR 3464, 8μm pore diameter) in DMEM. 100μL of wound-matched monocytes/macrophages (density of 372,000/mL in DMEM) were seeded in the transwell insert. Transwell cultures were incubated overnight at 37°C with 5% CO2.

Following incubation, transwells were removed and stained in crystal violet (Sigma-Aldrich V5265-500ML, 0.2% in methanol) for 10 minutes. Transwells were rinsed in water, and the inside was gently cleaned with a cotton swab. Inserts were left to dry for at least 24 hours. For the BMDM migration assay, 5-8 representative images from each transwell were taken for quantification. For the SCA1+/wound monocyte-macrophage assay, 20x tile images of each edge of the membrane as well as a region from the membrane center were taken for each insert. For both experiments, the number of migrated cells was calculated using QuPath software v0.3.2 (Bankhead et al., 2017). QuPath’s pixel classifier tool was used to train the software to recognize areas of positive crystal violet staining, which were then quantified per image. For the BMDM migration assay, data was reported as the average number of BMDMs per field of view for each transwell. For the SCA1+/wound monocyte-macrophage assay, the area analyzed was the same for each transwell, and the total number of migrated cells was reported.

### In situ hybridization for RNA detection (RNAscope)

Non-wounded skin and 1.5d wounds with 2 mm of non-wound skin cut around the edge from male C57 mice were sectioned within one week of harvest. Tissue sections were stained using the RNAscope^TM^ HiPlex v2 Assay kit (ACD Bio 324410) according to manufacturer instructions. Briefly, samples were fixed in 10% Neutral Buffered Formalin (NBF), dehydrated in ethanol, and treated with a protease from the kit (322340). Sections were treated with probes targeting *Pdgfra* (480661-T2) and *Ccl7* (446821-T3) or *Pdgfra* (480661-T5) and *Il33* (400591-T6), followed by kit amplifiers and T-fluorophores. As with immunofluorescence staining, slides were mounted in ProLong™ Gold Antifade mountant with DAPI.

### Immunofluorescence Staining

Naïve mouse back skin or wound beds were dissected, embedded in Scigen Tissue-Plus Optimal Cutting Temperature (OCT) Compound (Fisher 23-730-571), frozen, and cryosectioned into 14-16 µm sections on a Micron HM305 E Cryostat. Wounds were sectioned in their entirety and sequentially distributed across multiple slides to enable identification of the wound center and multi-parameter analysis of the same wound, as previously detailed (Haas et al., 2021). Frozen tissue sections were fixed in 4% formaldehyde and immunostained with the following antibodies: CD31 (BD Pharmingen, 1:50), ER-TR7 (Abcam, 1:500), F4/80 (Abcam, 1:200), ITGA6 (R&D Systems, 1:300), and pH3 (Abcam, 1:500) Slides were mounted in ProLong™ Gold Antifade mountant with DAPI (Invitrogen P36935) for nuclear staining. Additional information on antibodies used for immunofluorescence can be found in Supplementary Table S1.

### Imaging

All imaging was performed using a Zeiss AxioImager M2 (Zeiss, Oberkochen, Germany). Brightfield images of the crystal violet-stained transwells were taken with an Axiocam 305 color camera (Zeiss). Images of immunofluorescence or RNA *in situ* hybridization were acquired with an Orca camera (Hamamatsu, Hamamatsu, Japan). Tilescans of wound beds and the wound periphery were acquired using ZEN v3.0 software (Zeiss). For wounds, the two or three centralmost sections were imaged.

### Image analysis

Analyses and quantification were performed on the 2-3 most central sections of the wound. For analyses of wound beds, values from the different sections were averaged as a single wound bed value. Results were reported as single wounds or wound edges as described in figure legends and previously described (Schmidt and Horsley, 2013, Shook et al., 2020, Shook et al., 2018). For all analyses, wounds were taken from at least 3 different mice per group.

For F4/80 analysis in the wound center at 1.5dpw, nuclear signals associated with F4/80 staining were hand counted, and the wound bed area was quantified in Adobe Photoshop. For F4/80 analysis at the wound periphery, wound edges were cropped to 800μm in width from the edge of the wound and normalized to the same height in pixels. For consistency, 150-200 pixels of fascia below the panniculus carnosus were included in each image. Wound edge images were excluded if they differed significantly in the orientation of the skin layers compared to the rest of the set. F4/80+ cells in these images were identified using MATLAB 2023b. Using the regionprops() function, areas of staining in the images were called F4/80+ cells if there was at least 75% overlap of the nuclear periphery and F4/80 signals. For spatial analysis, images were divided into 20 bins along the superficial-deep and proximal-distal axes, and the number of cells in each bin was calculated. Total F4/80+ cells in each edge and their pixel coordinates on the image are also reported. To generate heat maps of F4/80 spatial distribution at the wound edge, the pixel coordinates from the cells in each image from a group were graphed as a rasterized, two-dimensional density plot in R v4.3.1. All wound edge images were analyzed individually and not averaged per wound.

For RNAscope analysis, UW images were cropped to 1.2 mm in length. 1.5d wounds were analyzed at the wound periphery and cropped to 1.2 mm from the wound edge with the same height. Images were analyzed using QuPath v0.3.2 (Bankhead et al., 2017). The CellDetection function was used to identify cells based on DAPI staining, followed by the SubcellularDetection function to identify cells with positive staining for *Pdgfra*, *Ccl7*, or *Il33*. The pixel coordinates of *Pdgfra*+*Ccl7*+ or *Pdgfra*+*Il33*+ on the images were subsequently obtained. Spatial heat maps were generated as described above in R v4.3.1.

To calculate re-epithelialization, ImageJ (FIJI) was used to measure the total wound width and the length not covered by ITGA6 as previously described (Yan et al., 2022). Wounds with continuous ITGA6+ epithelium were defined as closed and wounds with discontinuous ITGA6+ epithelium were defined as open.

The Color Range feature of Adobe Photoshop was used to identify CD31+ pixels, ER-TR7+ pixels, and total wound bed pixels to determine percent revascularization and fibroblast infiltration, respectively. Spatial analysis of CD31 was performed using ImageJ (FIJI) after cropping and blacking out non-wound areas in Photoshop. Proliferation analysis of pH3 staining was performed by identifying pH3+ cells in ImageJ (FIJI) and is reported as a function of the number of pH3+ cells in the region of interest divided by the total area of that region in mm^2^.

### Statistics

All statistics were performed using GraphPad Prism Version 9. Unpaired, two-tailed Student t-tests were used for comparisons between two groups, and one-way ANOVAs with Tukey’s Multiple Comparisons test for comparisons between multiple groups. For all statistical tests, *p* < 0.05 was considered significant.

## DATA AVAILABILITY STATEMENT

Single nuclei RNA-sequencing data generated in this study have been deposited in NCBI Gene Expression Omnibus (GEO) under accession number GSE265996.

## ACKNOWLEDGEMENTS

This work is supported in part by a start-up fund from The George Washington University to B.A.S, an NIH grant to B.A.S from NIAMS (AR082417), and by the National Science Foundation (2109688) to A.R. We thank members of the Shook and Rahnavard Labs for their assistance in processing samples and critical reading of the manuscript. snRNA-seq data are available at GEO: GSE265996 (reviewer token: ovwriiaqdlkztur).

## AUTHOR CONTRIBUTIONS

Conceptualization, V.M.A., M.R.H., P.O.C., R.C., A.R., B.A.S.; Methodology, V.M.A., P.O.C., R.C., C.G., M.F.M., A.R., B.A.S.; Formal Analysis, V.M.A., M.R.H., R.C., S.H., S.S., V.J., K.K., A.H., B.A.S.; Investigation, V.M.A., M.R.H., R.C., S.H., S.S., A.R.; Resources, A.R., B.N, B.A.S.; Writing – Original Draft, V.M.A., P.O.C., R.C., C.G., A.R., B.A.S.; Writing – Review & Editing, V.M.A., R.C., A.R., B.A.S.; Supervision, A.R., V.M.A., B.A.S.; Funding Acquisition, A.R., B.A.S.

## REFERENCES

Abbasi S, Sinha S, Labit E, Rosin NL, Yoon G, Rahmani W, et al. Distinct Regulatory Programs Control the Latent Regenerative Potential of Dermal Fibroblasts during Wound Healing. Cell Stem Cell 2020;27(3):396–412.e6.

Ahlers JMD, Falckenhayn C, Holzscheck N, Solé-Boldo L, Schütz S, Wenck H, et al. Single-Cell RNA Profiling of Human Skin Reveals Age-Related Loss of Dermal Sheath Cells and Their Contribution to a Juvenile Phenotype. Front Genet 2021;12:797747.

Al-Rikabi AHA, Tobin DJ, Riches-Suman K, Thornton MJ. Dermal fibroblasts cultured from donors with type 2 diabetes mellitus retain an epigenetic memory associated with poor wound healing responses. Sci Rep 2021;11(1):1474.

Atiakshin D, Kostin A, Trotsenko I, Samoilova V, Buchwalow I, Tiemann M. Carboxypeptidase A3-A Key Component of the Protease Phenotype of Mast Cells. Cells 2022;11(3).

Audu CO, Melvin WJ, Joshi AD, Wolf SJ, Moon JY, Davis FM, et al. Macrophage-specific inhibition of the histone demethylase JMJD3 decreases STING and pathologic inflammation in diabetic wound repair. Cell Mol Immunol 2022;19(11):1251–62.

Bankhead P, Loughrey MB, Fernández JA, Dombrowski Y, McArt DG, Dunne PD, et al. QuPath: Open source software for digital pathology image analysis. Sci Rep 2017;7(1):16878.

Bhattacharyya S, Wang W, Qin W, Cheng K, Coulup S, Chavez S, et al. TLR4-dependent fibroblast activation drives persistent organ fibrosis in skin and lung. JCI Insight 2018;3(13):e98850.

Boniakowski AE, Kimball AS, Joshi A, Schaller M, Davis FM, denDekker A, et al. Murine macrophage chemokine receptor CCR2 plays a crucial role in macrophage recruitment and regulated inflammation in wound healing. Eur J Immunol 2018;48(9):1445–55.

Boyd DF, Allen EK, Randolph AG, Guo X-zJ, Weng Y, Sanders CJ, et al. Exuberant fibroblast activity compromises lung function via ADAMTS4. Nature 2020;587(7834):466–71.

Brazil JC, Quiros M, Nusrat A, Parkos CA. Innate immune cell-epithelial crosstalk during wound repair. J Clin Invest 2019;129(8):2983–93.

Burke JP, Cunningham MF, Watson RW, Docherty NG, Coffey JC, O’Connell PR. Bacterial lipopolysaccharide promotes profibrotic activation of intestinal fibroblasts. Br J Surg 2010;97(7):1126–34.

Castanheira F, de Lima KA, Cebinelli GCM, Sônego F, Kanashiro A, Colon DF, et al. CCR5-Positive Inflammatory Monocytes are Crucial for Control of Sepsis. Shock 2019;52(5):e100–e6.

Chen C, Tang Y, Zhu X, Yang J, Liu Z, Chen Y, et al. P311 Promotes IL-4 Receptor‒Mediated M2 Polarization of Macrophages to Enhance Angiogenesis for Efficient Skin Wound Healing. Journal of Investigative Dermatology 2023;143(4):648–60.e6.

Chen G, Bracamonte-Baran W, Diny NL, Hou X, Talor MV, Fu K, et al. Sca-1(+) cardiac fibroblasts promote development of heart failure. Eur J Immunol 2018;48(9):1522–38.

Chen L, Mirza R, Kwon Y, DiPietro LA, Koh TJ. The murine excisional wound model: Contraction revisited. Wound Repair Regen 2015;23(6):874–7.

Choi ES, Jakubzick C, Carpenter KJ, Kunkel SL, Evanoff H, Martinez FJ, et al. Enhanced monocyte chemoattractant protein-3/CC chemokine ligand-7 in usual interstitial pneumonia. Am J Respir Crit Care Med 2004;170(5):508–15.

Chung MI, Bujnis M, Barkauskas CE, Kobayashi Y, Hogan BLM. Niche-mediated BMP/SMAD signaling regulates lung alveolar stem cell proliferation and differentiation. Development 2018;145(9).

Correa-Gallegos D, Ye H, Dasgupta B, Sardogan A, Kadri S, Kandi R, et al. CD201(+) fascia progenitors choreograph injury repair. Nature 2023.

Crane MJ, Daley JM, van Houtte O, Brancato SK, Henry WL, Jr., Albina JE. The monocyte to macrophage transition in the murine sterile wound. PLoS One 2014;9(1):e86660.

Daley JM, Brancato SK, Thomay AA, Reichner JS, Albina JE. The phenotype of murine wound macrophages. J Leukoc Biol 2010;87(1):59–67.

Dave RK, Hume DA, Elsegood C, Kellie S. CD148/DEP-1 association with areas of cytoskeletal organisation in macrophages. Experimental Cell Research 2009;315(10):1734–44.

De Filippo K, Dudeck A, Hasenberg M, Nye E, van Rooijen N, Hartmann K, et al. Mast cell and macrophage chemokines CXCL1/CXCL2 control the early stage of neutrophil recruitment during tissue inflammation. Blood 2013;121(24):4930–7.

de Oliveira S, Rosowski EE, Huttenlocher A. Neutrophil migration in infection and wound repair: going forward in reverse. Nat Rev Immunol 2016;16(6):378–91.

Dewald O, Zymek P, Winkelmann K, Koerting A, Ren G, Abou-Khamis T, et al. CCL2/Monocyte Chemoattractant Protein-1 regulates inflammatory responses critical to healing myocardial infarcts. Circ Res 2005;96(8):881–9.

Driskell RR, Lichtenberger BM, Hoste E, Kretzschmar K, Simons BD, Charalambous M, et al. Distinct fibroblast lineages determine dermal architecture in skin development and repair. Nature 2013;504(7479):277–81.

Eming SA, Martin P, Tomic-Canic M. Wound repair and regeneration: mechanisms, signaling, and translation. Sci Transl Med 2014;6(265):265sr6.

Farbehi N, Patrick R, Dorison A, Xaymardan M, Janbandhu V, Wystub-Lis K, et al. Single-cell expression profiling reveals dynamic flux of cardiac stromal, vascular and immune cells in health and injury. Elife 2019;8.

Ferrer RA, Saalbach A, Grünwedel M, Lohmann N, Forstreuter I, Saupe S, et al. Dermal Fibroblasts Promote Alternative Macrophage Activation Improving Impaired Wound Healing. Journal of Investigative Dermatology 2017;137(4):941–50.

Foster DS, Januszyk M, Yost KE, Chinta MS, Gulati GS, Nguyen AT, et al. Integrated spatial multiomics reveals fibroblast fate during tissue repair. Proc Natl Acad Sci U S A 2021;118(41).

Frech S, Forsthuber A, Korosec A, Lipp K, Kozumov V, Lichtenberger BM. Hedgehog Signaling in Papillary Fibroblasts Is Essential for Hair Follicle Regeneration during Wound Healing. Journal of Investigative Dermatology 2022;142(6):1737–48.e5.

Fujiwara H, Ferreira M, Donati G, Marciano DK, Linton JM, Sato Y, et al. The Basement Membrane of Hair Follicle Stem Cells Is a Muscle Cell Niche. Cell 2011;144(4):577–89.

Goren I, Allmann N, Yogev N, Schürmann C, Linke A, Holdener M, et al. A transgenic mouse model of inducible macrophage depletion: effects of diphtheria toxin-driven lysozyme M-specific cell lineage ablation on wound inflammatory, angiogenic, and contractive processes. Am J Pathol 2009;175(1):132–47.

Gould L, Abadir P, Brem H, Carter M, Conner-Kerr T, Davidson J, et al. Chronic wound repair and healing in older adults: current status and future research. Wound Repair Regen 2015;23(1):1–13.

Guenin-Mace L, Konieczny P, Naik S. Immune-Epithelial Cross Talk in Regeneration and Repair. Annu Rev Immunol 2023;41:207–28.

Guerrero-Juarez CF, Dedhia PH, Jin S, Ruiz-Vega R, Ma D, Liu Y, et al. Single-cell analysis reveals fibroblast heterogeneity and myeloid-derived adipocyte progenitors in murine skin wounds. Nature communications 2019;10(1):650-.

Gurtner GC, Werner S, Barrandon Y, Longaker MT. Wound repair and regeneration. Nature 2008;453(7193):314–21.

Haas MR, Nguyen DV, Shook BA. Recovery of Altered Diabetic Myofibroblast Heterogeneity and Gene Expression Are Associated with CD301b+ Macrophages. Biomedicines 2021.

Habib N, Avraham-Davidi I, Basu A, Burks T, Shekhar K, Hofree M, et al. Massively parallel single-nucleus RNA-seq with DroNc-seq. Nature Methods 2017;14(10):955–8.

Hafemeister C, Satija R. Normalization and variance stabilization of single-cell RNA-seq data using regularized negative binomial regression. Genome Biology 2019;20(1):296.

Hao Y, Hao S, Andersen-Nissen E, Mauck WM, Zheng S, Butler A, et al. Integrated analysis of multimodal single-cell data. Cell 2021;184(13):3573–87.e29.

He C-c, Song T-c, Qi R-q, Gao X-H. Integrated single-cell and spatial transcriptomics reveals heterogeneity of fibroblast and pivotal genes in psoriasis. Scientific Reports 2023;13(1):17134.

He R, Yin H, Yuan B, Liu T, Luo L, Huang P, et al. IL-33 improves wound healing through enhanced M2 macrophage polarization in diabetic mice. Mol Immunol 2017;90:42–9.

Hoffman EP, Brown RH, Jr., Kunkel LM. Dystrophin: the protein product of the Duchenne muscular dystrophy locus. Cell 1987;51(6):919–28.

Hu KH, Kuhn NF, Courau T, Tsui J, Samad B, Ha P, et al. Transcriptional space-time mapping identifies concerted immune and stromal cell patterns and gene programs in wound healing and cancer. Cell Stem Cell 2023;30(6):885–903.e10.

Janson DG, Saintigny G, van Adrichem A, Mahé C, El Ghalbzouri A. Different gene expression patterns in human papillary and reticular fibroblasts. J Invest Dermatol 2012;132(11):2565–72.

Januszyk M, Chen K, Henn D, Foster DS, Borrelli MR, Bonham CA, et al. Characterization of Diabetic and Non-Diabetic Foot Ulcers Using Single-Cell RNA-Sequencing. Micromachines (Basel) 2020;11(9).

Jiang D, Christ S, Correa-Gallegos D, Ramesh P, Kalgudde Gopal S, Wannemacher J, et al. Injury triggers fascia fibroblast collective cell migration to drive scar formation through N-cadherin. Nature Communications 2020;11(1):5653.

Jo HY, Seo HH, Gil D, Park Y, Han HJ, Han HW, et al. Single-Cell RNA Sequencing of Human Pluripotent Stem Cell-Derived Macrophages for Quality Control of The Cell Therapy Product. Front Genet 2021;12:658862.

Joost S, Annusver K, Jacob T, Sun X, Dalessandri T, Sivan U, et al. The Molecular Anatomy of Mouse Skin during Hair Growth and Rest. Cell Stem Cell 2020;26(3):441–57.e7.

Joshi N, Pohlmeier L, Ben-Yehuda Greenwald M, Haertel E, Hiebert P, Kopf M, et al. Comprehensive characterization of myeloid cells during wound healing in healthy and healing-impaired diabetic mice. Eur J Immunol 2020;50(9):1335–49.

Kawaguchi M, Takahashi M, Hata T, Kashima Y, Usui F, Morimoto H, et al. Inflammasome activation of cardiac fibroblasts is essential for myocardial ischemia/reperfusion injury. Circulation 2011;123(6):594–604.

Kim Y-J, Cho SY, Yun CH, Moon YS, Lee TR, Kim SH. Transcriptional activation of Cidec by PPARγ2 in adipocyte. Biochemical and Biophysical Research Communications 2008;377(1):297–302.

Kitanaka N, Nakano R, Sugiura K, Kitanaka T, Namba S, Konno T, et al. Interleukin-1β promotes interleulin-6 expression via ERK1/2 signaling pathway in canine dermal fibroblasts. PLoS One 2019;14(7):e0220262.

Konger RL, Derr-Yellin E, Zimmers TA, Katona T, Xuei X, Liu Y, et al. Epidermal PPARγ Is a Key Homeostatic Regulator of Cutaneous Inflammation and Barrier Function in Mouse Skin. International Journal of Molecular Sciences 2021;22(16):8634.

Korosec A, Frech S, Gesslbauer B, Vierhapper M, Radtke C, Petzelbauer P, et al. Lineage Identity and Location within the Dermis Determine the Function of Papillary and Reticular Fibroblasts in Human Skin. J Invest Dermatol 2019;139(2):342–51.

Krämer A, Green J, Pollard J, Jr., Tugendreich S. Causal analysis approaches in Ingenuity Pathway Analysis. Bioinformatics 2014;30(4):523–30.

Krausgruber T, Fortelny N, Fife-Gernedl V, Senekowitsch M, Schuster LC, Lercher A, et al. Structural cells are key regulators of organ-specific immune responses. Nature 2020;583(7815):296–302.

Le Fournis C, Jeanneau C, Giraud T, El Karim I, Lundy FT, About I. Fibroblasts Control Macrophage Differentiation during Pulp Inflammation. Journal of Endodontics 2021;47(9):1427–34.

Leavitt T, Hu MS, Borrelli MR, Januszyk M, Garcia JT, Ransom RC, et al. Prrx1 Fibroblasts Represent a Pro-fibrotic Lineage in the Mouse Ventral Dermis. Cell Rep 2020;33(6):108356.

Lebre MC, van der Aar AM, van Baarsen L, van Capel TM, Schuitemaker JH, Kapsenberg ML, et al. Human keratinocytes express functional Toll-like receptor 3, 4, 5, and 9. J Invest Dermatol 2007;127(2):331–41.

Li H, Humphreys BD. Mouse kidney nuclear isolation and library preparation for single-cell combinatorial indexing RNA sequencing. STAR Protoc 2022;3(4):101904.

Lichtenberger BM, Mastrogiannaki M, Watt FM. Epidermal β-catenin activation remodels the dermis via paracrine signalling to distinct fibroblast lineages. Nature communications 2016;7:10537-.

Lorenz DR, Misra V, Gabuzda D. Transcriptomic analysis of monocytes from HIV-positive men on antiretroviral therapy reveals effects of tobacco smoking on interferon and stress response systems associated with depressive symptoms. Hum Genomics 2019;13(1):59.

Lu H, Huang D, Ransohoff RM, Zhou L. Acute skeletal muscle injury: CCL2 expression by both monocytes and injured muscle is required for repair. Faseb j 2011;25(10):3344–55.

Lucas T, Waisman A, Ranjan R, Roes J, Krieg T, Müller W, et al. Differential roles of macrophages in diverse phases of skin repair. J Immunol 2010;184(7):3964–77.

Luecken MD, Theis FJ. Current best practices in single-cell RNA-seq analysis: a tutorial. Molecular Systems Biology 2019;15(6):e8746.

Mahmoudi S, Mancini E, Xu L, Moore A, Jahanbani F, Hebestreit K, et al. Heterogeneity in old fibroblasts is linked to variability in reprogramming and wound healing. Nature 2019;574(7779):553–8.

Mallick H, Chatterjee S, Chowdhury S, Chatterjee S, Rahnavard A, Hicks SC. Differential expression of single-cell RNA-seq data using Tweedie models. Stat Med 2022;41(18):3492–510.

Maraux M, Gaillardet A, Gally A, Saas P, Cherrier T. Human primary neutrophil mRNA does not contaminate human resolving macrophage mRNA after efferocytosis. J Immunol Methods 2020;483:112810.

Mirza R, DiPietro LA, Koh TJ. Selective and specific macrophage ablation is detrimental to wound healing in mice. Am J Pathol 2009;175(6):2454–62.

Mirza R, Koh TJ. Dysregulation of monocyte/macrophage phenotype in wounds of diabetic mice. Cytokine 2011;56(2):256–64.

Mirza RE, Fang MM, Weinheimer-Haus EM, Ennis WJ, Koh TJ. Sustained inflammasome activity in macrophages impairs wound healing in type 2 diabetic humans and mice. Diabetes 2014;63(3):1103–14.

Mouton AJ, Ma Y, Rivera Gonzalez OJ, Daseke MJ, 2nd, Flynn ER, Freeman TC, et al. Fibroblast polarization over the myocardial infarction time continuum shifts roles from inflammation to angiogenesis. Basic Res Cardiol 2019;114(2):6-.

Oh S, Lee JH, Kim HM, Batsukh S, Sung MJ, Lim TH, et al. Poly-L-Lactic Acid Fillers Improved Dermal Collagen Synthesis by Modulating M2 Macrophage Polarization in Aged Animal Skin. Cells 2023;12(9).

Oppermann M. Chemokine receptor CCR5: insights into structure, function, and regulation. Cell Signal 2004;16(11):1201–10.

Osborne JM, den Elzen N, Lichanska AM, Costelloe EO, Yamada T, Cassady AI, et al. Murine DEP-1, a receptor protein tyrosine phosphatase, is expressed in macrophages and is regulated by CSF-1 and LPS. J Leukoc Biol 1998;64(5):692–701.

Osorio D, Cai JJ. Systematic determination of the mitochondrial proportion in human and mice tissues for single-cell RNA-sequencing data quality control. Bioinformatics 2021;37(7):963–7.

Paish HL, Kalson NS, Smith GR, Del Carpio Pons A, Baldock TE, Smith N, et al. Fibroblasts Promote Inflammation and Pain via IL-1α Induction of the Monocyte Chemoattractant Chemokine (C-C Motif) Ligand 2. Am J Pathol 2018;188(3):696–714.

Pang J, Maienschein-Cline M, Koh TJ. Enhanced Proliferation of Ly6C+ Monocytes/Macrophages Contributes to Chronic Inflammation in Skin Wounds of Diabetic Mice. The Journal of Immunology 2021;206(3):621–30.

Pang J, Maienschein-Cline M, Koh TJ. Monocyte/Macrophage Heterogeneity during Skin Wound Healing in Mice. The Journal of Immunology 2022;209(10):1999–2011.

Pappritz K, Savvatis K, Koschel A, Miteva K, Tschöpe C, Van Linthout S. Cardiac (myo)fibroblasts modulate the migration of monocyte subsets. Scientific Reports 2018;8(1):5575.

Parfejevs V, Debbache J, Shakhova O, Schaefer SM, Glausch M, Wegner M, et al. Injury-activated glial cells promote wound healing of the adult skin in mice. Nat Commun 2018;9(1):236.

Phan QM, Fine GM, Salz L, Herrera GG, Wildman B, Driskell IM, et al. Lef1 expression in fibroblasts maintains developmental potential in adult skin to regenerate wounds. Elife 2020;9.

Philippeos C, Telerman SB, Oulès B, Pisco AO, Shaw TJ, Elgueta R, et al. Spatial and Single-Cell Transcriptional Profiling Identifies Functionally Distinct Human Dermal Fibroblast Subpopulations. J Invest Dermatol 2018;138(4):811–25.

Ploeger DT, Hosper NA, Schipper M, Koerts JA, de Rond S, Bank RA. Cell plasticity in wound healing: paracrine factors of M1/ M2 polarized macrophages influence the phenotypical state of dermal fibroblasts. Cell Commun Signal 2013;11(1):29.

Puzzi L, Borin D, Martinelli V, Mestroni L, Kelsell DP, Sbaizero O. Cellular biomechanics impairment in keratinocytes is associated with a C-terminal truncated desmoplakin: An atomic force microscopy investigation. Micron 2018;106:27–33.

Qu J, Tanis SEJ, Smits JPH, Kouwenhoven EN, Oti M, van den Bogaard EH, et al. Mutant p63 Affects Epidermal Cell Identity through Rewiring the Enhancer Landscape. Cell Rep 2018;25(12):3490–503.e4.

Ridiandries A, Tan JTM, Bursill CA. The Role of Chemokines in Wound Healing. Int J Mol Sci 2018;19(10).

Rinkevich Y, Walmsley GG, Hu MS, Maan ZN, Newman AM, Drukker M, et al. Skin fibrosis. Identification and isolation of a dermal lineage with intrinsic fibrogenic potential. Science 2015;348(6232).

Rittié L, Tejasvi T, Harms PW, Xing X, Nair RP, Gudjonsson JE, et al. Sebaceous Gland Atrophy in Psoriasis: An Explanation for Psoriatic Alopecia? Journal of Investigative Dermatology 2016;136(9):1792–800.

Rognoni E, Gomez C, Pisco AO, Rawlins EL, Simons BD, Watt FM, et al. Inhibition of β-catenin signalling in dermal fibroblasts enhances hair follicle regeneration during wound healing. Development 2016;143(14):2522–35.

Rosen ED, Sarraf P, Troy AE, Bradwin G, Moore K, Milstone DS, et al. PPARγ Is Required for the Differentiation of Adipose Tissue In Vivo and In Vitro. Molecular Cell 1999;4(4):611–7.

Roupé KM, Nybo M, Sjöbring U, Alberius P, Schmidtchen A, Sørensen OE. Injury is a major inducer of epidermal innate immune responses during wound healing. J Invest Dermatol 2010;130(4):1167–77.

Satija R, Farrell JA, Gennert D, Schier AF, Regev A. Spatial reconstruction of single-cell gene expression data. Nat Biotechnol 2015;33(5):495–502.

Sawant KV, Sepuru KM, Lowry E, Penaranda B, Frevert CW, Garofalo RP, et al. Neutrophil recruitment by chemokines Cxcl1/KC and Cxcl2/MIP2: Role of Cxcr2 activation and glycosaminoglycan interactions. J Leukoc Biol 2021;109(4):777–91.

Sawaya AP, Stone RC, Brooks SR, Pastar I, Jozic I, Hasneen K, et al. Deregulated immune cell recruitment orchestrated by FOXM1 impairs human diabetic wound healing. Nat Commun 2020;11(1):4678.

Schmidt BA, Horsley V. Intradermal adipocytes mediate fibroblast recruitment during skin wound healing. Development 2013;140(7):1517–27.

Shi C, Jia T, Mendez-Ferrer S, Hohl TM, Serbina NV, Lipuma L, et al. Bone marrow mesenchymal stem and progenitor cells induce monocyte emigration in response to circulating toll-like receptor ligands. Immunity 2011;34(4):590–601.

Shook B, Xiao E, Kumamoto Y, Iwasaki A, Horsley V. CD301b+ Macrophages Are Essential for Effective Skin Wound Healing. J Invest Dermatol 2016;136(9):1885–91.

Shook BA, Wasko RR, Mano O, Rutenberg-Schoenberg M, Rudolph MC, Zirak B, et al. Dermal Adipocyte Lipolysis and Myofibroblast Conversion Are Required for Efficient Skin Repair. Cell Stem Cell 2020;26(6):880–95.e6.

Shook BA, Wasko RR, Rivera-Gonzalez GC, Salazar-Gatzimas E, López-Giráldez F, Dash BC, et al. Myofibroblast proliferation and heterogeneity are supported by macrophages during skin repair. Science 2018;362(6417).

Siddhuraj P, Clausson CM, Sanden C, Alyamani M, Kadivar M, Marsal J, et al. Lung Mast Cells Have a High Constitutive Expression of Carboxypeptidase A3 mRNA That Is Independent from Granule-Stored CPA3. Cells 2021;10(2).

Sinha S, Sparks HD, Labit E, Robbins HN, Gowing K, Jaffer A, et al. Fibroblast inflammatory priming determines regenerative versus fibrotic skin repair in reindeer. Cell 2022;185(25):4717–36.e25.

Spindler V, Heupel WM, Efthymiadis A, Schmidt E, Eming R, Rankl C, et al. Desmocollin 3-mediated binding is crucial for keratinocyte cohesion and is impaired in pemphigus. J Biol Chem 2009;284(44):30556–64.

Stuart T, Butler A, Hoffman P, Hafemeister C, Papalexi E, Mauck WM, III, et al. Comprehensive Integration of Single-Cell Data. Cell 2019;177(7):1888–902.e21.

Sundberg JP, Shen T, Fiehn O, Rice RH, Silva KA, Kennedy VE, et al. Sebaceous gland abnormalities in fatty acyl CoA reductase 2 (Far2) null mice result in primary cicatricial alopecia. PLoS One 2018;13(10):e0205775.

Suresh MV, Yu B, Machado-Aranda D, Bender MD, Ochoa-Frongia L, Helinski JD, et al. Role of macrophage chemoattractant protein-1 in acute inflammation after lung contusion. Am J Respir Cell Mol Biol 2012;46(6):797–806.

Tombor LS, John D, Glaser SF, Luxán G, Forte E, Furtado M, et al. Single cell sequencing reveals endothelial plasticity with transient mesenchymal activation after myocardial infarction. Nature Communications 2021;12(1):681.

Tsou C-L, Peters W, Si Y, Slaymaker S, Aslanian AM, Weisberg SP, et al. Critical roles for CCR2 and MCP-3 in monocyte mobilization from bone marrow and recruitment to inflammatory sites. The Journal of Clinical Investigation 2007;117(4):902–9.

Villarreal-Ponce A, Tiruneh MW, Lee J, Guerrero-Juarez CF, Kuhn J, David JA, et al. Keratinocyte-Macrophage Crosstalk by the Nrf2/Ccl2/EGF Signaling Axis Orchestrates Tissue Repair. Cell Rep 2020;33(8):108417.

Vu R, Jin S, Sun P, Haensel D, Nguyen QH, Dragan M, et al. Wound healing in aged skin exhibits systems-level alterations in cellular composition and cell-cell communication. Cell Reports 2022;40(5):111155.

Wall IB, Moseley R, Baird DM, Kipling D, Giles P, Laffafian I, et al. Fibroblast dysfunction is a key factor in the non-healing of chronic venous leg ulcers. J Invest Dermatol 2008;128(10):2526–40.

Weischenfeldt J, Porse B. Bone Marrow-Derived Macrophages (BMM): Isolation and Applications. CSH Protoc 2008;2008:pdb.prot5080.

Wigerblad G, Cao Q, Brooks S, Naz F, Gadkari M, Jiang K, et al. Single-Cell Analysis Reveals the Range of Transcriptional States of Circulating Human Neutrophils. J Immunol 2022;209(4):772–82.

Willenborg S, Lucas T, van Loo G, Knipper JA, Krieg T, Haase I, et al. CCR2 recruits an inflammatory macrophage subpopulation critical for angiogenesis in tissue repair. Blood 2012;120(3):613–25.

Williams DW, Greenwell-Wild T, Brenchley L, Dutzan N, Overmiller A, Sawaya AP, et al. Human oral mucosa cell atlas reveals a stromal-neutrophil axis regulating tissue immunity. Cell 2021;184(15):4090–104.e15.

Witowski J, Tayama H, Ksiazek K, Wanic-Kossowska M, Bender TO, Jörres A. Human peritoneal fibroblasts are a potent source of neutrophil-targeting cytokines: a key role of IL-1beta stimulation. Lab Invest 2009;89(4):414–24.

Wong VW, Rustad KC, Akaishi S, Sorkin M, Glotzbach JP, Januszyk M, et al. Focal adhesion kinase links mechanical force to skin fibrosis via inflammatory signaling. Nat Med 2011;18(1):148–52.

Wood S, Jayaraman V, Huelsmann EJ, Bonish B, Burgad D, Sivaramakrishnan G, et al. Pro-inflammatory chemokine CCL2 (MCP-1) promotes healing in diabetic wounds by restoring the macrophage response. PLoS One 2014;9(3):e91574.

Wu L, Ong S, Talor MV, Barin JG, Baldeviano GC, Kass DA, et al. Cardiac fibroblasts mediate IL-17A-driven inflammatory dilated cardiomyopathy. J Exp Med 2014;211(7):1449–64.

Xu C, Su Z. Identification of cell types from single-cell transcriptomes using a novel clustering method. Bioinformatics 2015;31(12):1974–80.

Yan S, Ripamonti R, Kawabe H, Ben-Yehuda Greenwald M, Werner S. NEDD4-1 Is a Key Regulator of Epidermal Homeostasis and Wound Repair. Journal of Investigative Dermatology 2022;142(6):1703–13.e11.

Yao C, Oh JH, Lee DH, Bae JS, Jin CL, Park CH, et al. Toll-like receptor family members in skin fibroblasts are functional and have a higher expression compared to skin keratinocytes. Int J Mol Med 2015;35(5):1443–50.

Yu M, Hu J, Zhu MX, Zhao T, Liang W, Wen S, et al. Cardiac fibroblasts recruit Th17 cells infiltration into myocardium by secreting CCL20 in CVB3-induced acute viral myocarditis. Cell Physiol Biochem 2013;32(5):1437–50.

Zhang J, Wu L, Qu JM, Bai CX, Merrilees MJ, Black PN. Pro-inflammatory phenotype of COPD fibroblasts not compatible with repair in COPD lung. J Cell Mol Med 2012;16(7):1522–32.

Zhang Z, Shao M, Hepler C, Zi Z, Zhao S, An YA, et al. Dermal adipose tissue has high plasticity and undergoes reversible dedifferentiation in mice. J Clin Invest 2019;129(12):5327–42.

Zheng GX, Terry JM, Belgrader P, Ryvkin P, Bent ZW, Wilson R, et al. Massively parallel digital transcriptional profiling of single cells. Nat Commun 2017;8:14049.

Zhou X, Franklin RA, Adler M, Carter TS, Condiff E, Adams TS, et al. Microenvironmental sensing by fibroblasts controls macrophage population size. Proc Natl Acad Sci U S A 2022;119(32):e2205360119.

Zhuang L, Zong X, Yang Q, Fan Q, Tao R. Interleukin-34-NF-κB signaling aggravates myocardial ischemic/reperfusion injury by facilitating macrophage recruitment and polarization. EBioMedicine 2023;95:104744.

