## Supplemental Data for "Deep skin fibroblast-mediated macrophage recruitment supports acute wound healing"

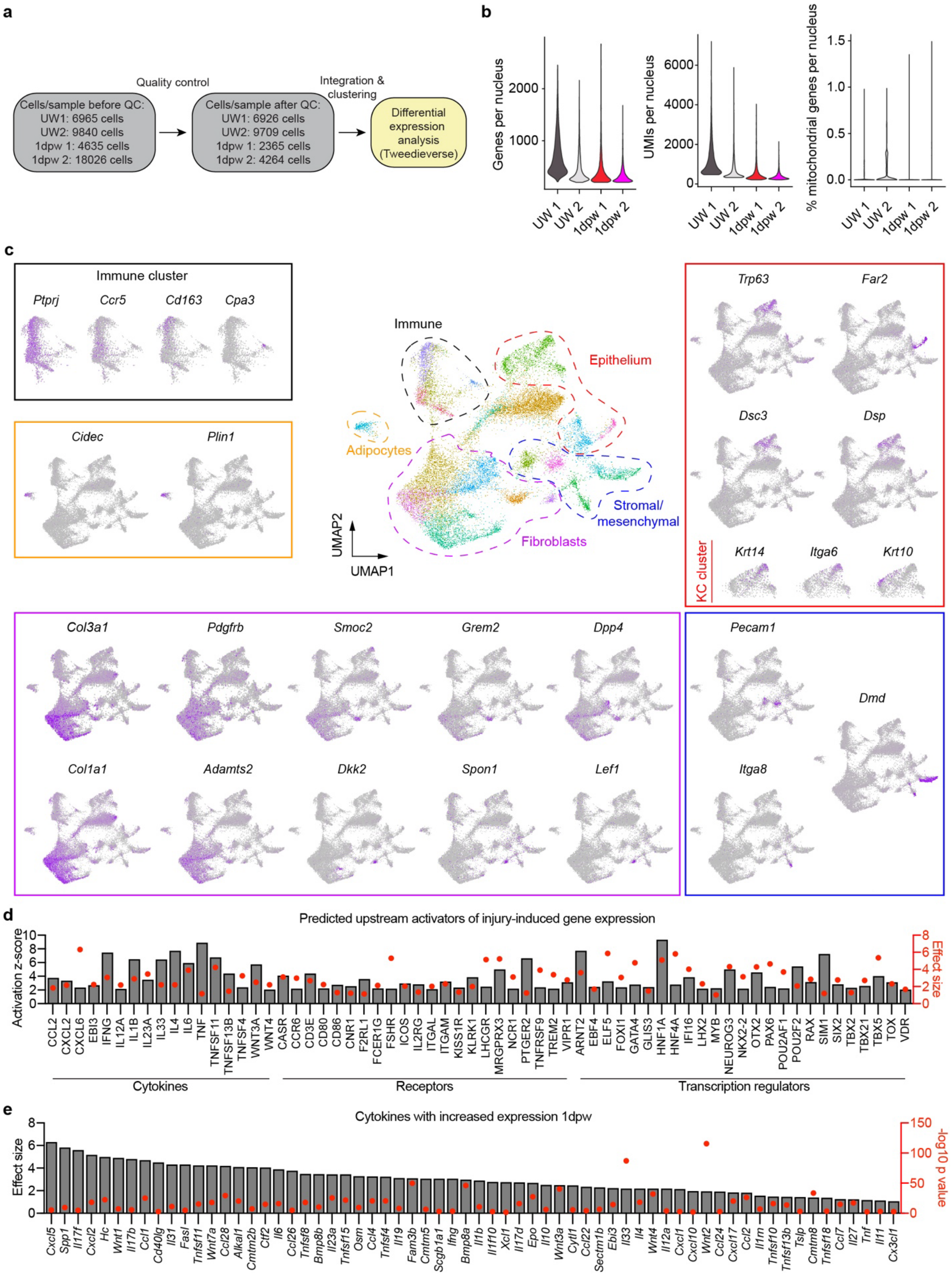
a
Quality control
Integration & clustering
Cells/sample before QC:
UW1: 6965 cells
UW2: 9840 cells
1dpw 1: 4635 cells
1dpw 2: 18026 cells
Cells/sample after QC:
UW1: 6926 cells
UW2: 9709 cells
1dpw 1: 2365 cells
1dpw 2: 4264 cells
Differential expression analysis (Tweedieverse)
b
Genes per nucleus
UMIs per nucleus
% mitochondrial genes per nucleus
UW 1
UW 2
1dpw 1
1dpw 2
c
Immune cluster
Ptprj
Ccr5
Cd163
Cpa3
Cidec
Plin1
Immune
Epithelium
Adipocytes
Stromal/ mesenchymal
Fibroblasts
UMAP1
UMAP2
Trp63
Far2
Dsc3
Dsp
KC cluster
Krt14
Itga6
Krt10
Col3a1
Pdgfrb
Smoc2
Grem2
Dpp4
Col1a1
Adamts2
Dkk2
Spon1
Lef1
Pecam1
Dmd
Itga8
d
Predicted upstream activators of injury-induced gene expression
Activation z-score
Effect size
CCL2 CXCL2 CXCL6 EBI3 IFNG IL12A IL1B IL23A IL33 IL4 IL6 TNF TNFSF11 TNFSF13B TNFSF4 WNT3A WNT4
CASR CCR6 CD3E CD80 CD86 CNR1 F2RL1 FCER1G FSHR ICOS IL2RG ITGAL ITGAM KISS1R KLRK1 LHCGR MRGPRX3 NCR1 PTGER2 TNFRSF9 TREM2 VIPR1
ARNT2 EBF4 ELF5 FOXI1 GATA4 GLIS3 HNF1A HNF4A IFI16 LHX2 MYB NEUROG3 NKX2-2 OTX2 PAX6 POU2AF1 POU2F2 RAX SIM1 SIX2 TBX2 TBX21 TBX5 TOX VDR
Cytokines
Receptors
Transcription regulators
e
Cytokines with increased expression 1dpw
Effect size
-log10 p value
Cxcl5 Spp1 Il17f Cxcl2 Hc Wnt1 Il17b Ccl1 Cd40lg Il31 Fasl Tnfsf11 Wnt7a Ccl28 Alkal1 Cmtm2b Ctf2 Il6 Ccl26 Tnfsf8 Bmp8b Il23a Tnfsf15 Osm Ccl4 Tnfsf4 Il19 Fam3b Cmtm5 Scgb1a1 Ifng Bmp8a Il1b Il1f10 Xcl1 Il17d Epo Il10 Wnt3a Cytl1 Ccl22 Sectm1b Ebi3 Il33 Il4 Wnt4 Il12a Cxcl1 Cxcl10 Wnt2 Ccl24 Cxcl17 Ccl2 Il1rn Tnfsf10 Tnfsf13b Tslp Cmtm8 Tnfsf18 Ccl7 Il27 Tnf Il11 Cx3cl1

**Supplementary Figure S1. Identification of cellular subsets and predicted regulators of injury-induced gene expression from single nuclei sequencing of skin.**

(a) Pipeline for snRNA-seq data analysis.

(b) Number of genes identified per nucleus (left), number of unique molecular identifiers (UMIs) per nucleus (center), and percentage of mitochondrial genes per nucleus (right) in each sample.

(c) Feature plots for genes associated with specific cell populations.

(d) Upstream activators of upregulated gene expression at 1dpw identified by Ingenuity Pathway Analysis.

(e) Cytokines with significant effect size at 1dpw.

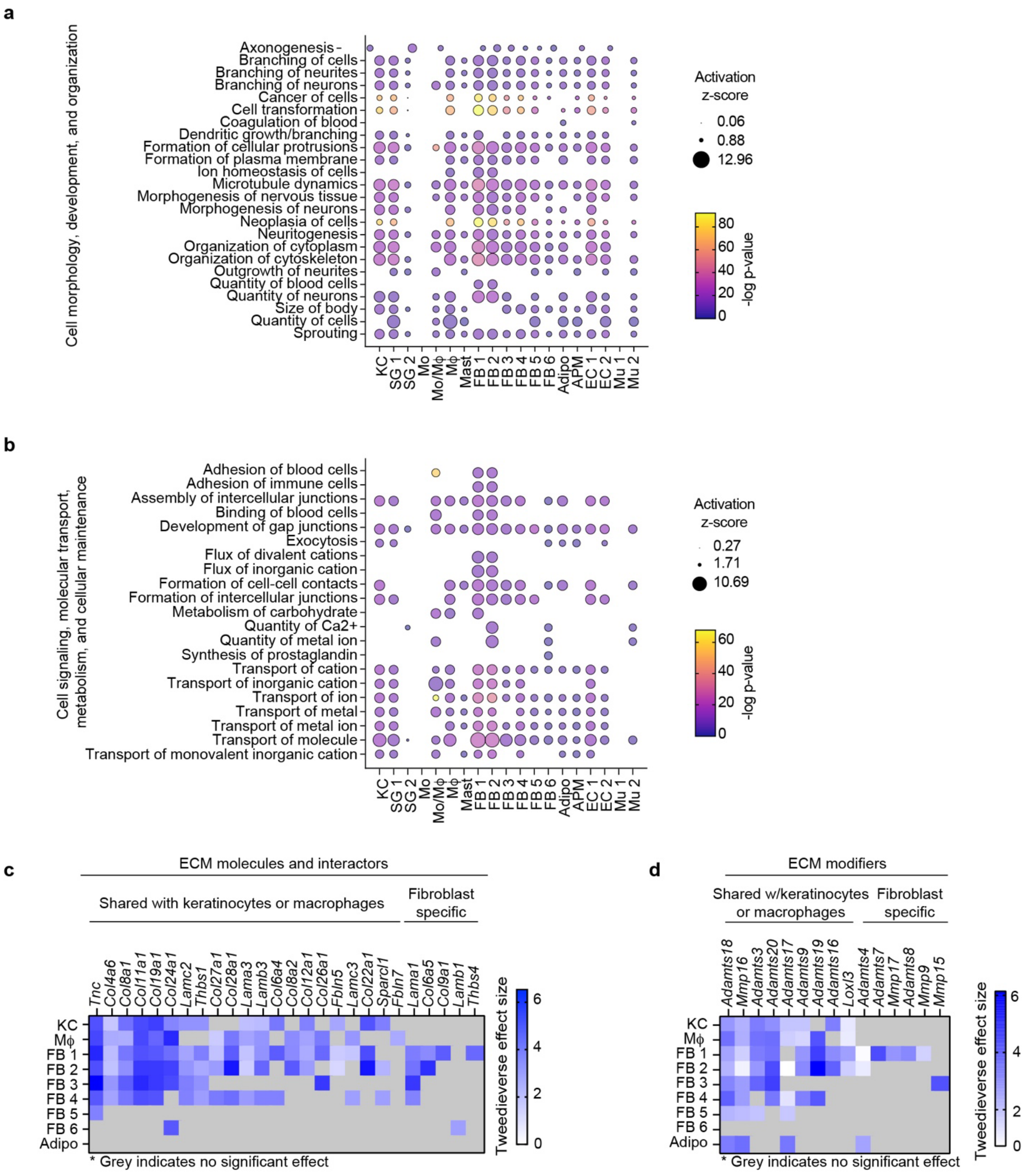
a
Cell morphology, development, and organization
Axonogenesis
Branching of cells
Branching of neurites
Branching of neurons
Cancer of cells
Cell transformation
Coagulation of blood
Dendritic growth/branching
Formation of cellular protrusions
Formation of plasma membrane
Ion homeostasis of cells
Microtubule dynamics
Morphogenesis of nervous tissue
Morphogenesis of neurons
Neoplasia of cells
Neuritogenesis
Organization of cytoplasm
Organization of cytoskeleton
Outgrowth of neurites
Quantity of blood cells
Quantity of neurons
Size of body
Quantity of cells
Sprouting
KC
SG 1
SG 2
Mo
Mo/Mϕ
Mϕ
Mast
FB 1
FB 2
FB 3
FB 4
FB 5
FB 6
Adipo
APM
EC 1
EC 2
Mu 1
Mu 2
Activation z-score
0.06
0.88
12.96
-log p-value
80
60
40
20
0
b
Cell signaling, molecular transport, metabolism, and cellular maintenance
Adhesion of blood cells
Adhesion of immune cells
Assembly of intercellular junctions
Binding of blood cells
Development of gap junctions
Exocytosis
Flux of divalent cations
Flux of inorganic cation
Formation of cell-cell contacts
Formation of intercellular junctions
Metabolism of carbohydrate
Quantity of Ca2+
Quantity of metal ion
Synthesis of prostaglandin
Transport of cation
Transport of inorganic cation
Transport of ion
Transport of metal
Transport of metal ion
Transport of molecule
Transport of monovalent inorganic cation
KC
SG 1
SG 2
Mo
Mo/Mϕ
Mϕ
Mast
FB 1
FB 2
FB 3
FB 4
FB 5
FB 6
Adipo
APM
EC 1
EC 2
Mu 1
Mu 2
Activation z-score
0.27
1.71
10.69
-log p-value
60
40
20
0
c
ECM molecules and interactors
Shared with keratinocytes or macrophages
Fibroblast specific
Tnc
Col4a6
Col8a1
Col11a1
Col19a1
Col24a1
Lamc2
Thbs1
Col27a1
Col28a1
Lama3
Lamb3
Col6a4
Col8a2
Col12a1
Col26a1
Fbln5
Lamc3
Col22a1
Sparcl1
Fbln7
Lama1
Col6a5
Col9a1
Lamb1
Thbs4
KC
Mϕ
FB 1
FB 2
FB 3
FB 4
FB 5
FB 6
Adipo
* Grey indicates no significant effect
Tweedieverse effect size
6
4
2
0
d
ECM modifiers
Shared w/keratinocytes or macrophages
Fibroblast specific
Adamts18
Mmp16
Adamts3
Adamts20
Adamts17
Adamts9
Adamts19
Adamts16
Loxl3
Adamts4
Adamts7
Mmp17
Adamts8
Mmp9
Mmp15
KC
Mϕ
FB 1
FB 2
FB 3
FB 4
FB 5
FB 6
Adipo
* Grey indicates no significant effect
Tweedieverse effect size
6
4
2
0

**Supplementary Figure S2. Injury-induced changes in gene expression generate distinct and redundant predicted biological functions in cellular subsets.**

(a-b) Bubble matrices of Biofunctions associated with upregulated gene expression in each cell cluster 1dpw. Matrices are divided by terms relating to cell morphology, development, and organization (a) or signaling, transport, metabolism, and cell maintenance (b).

(c-d) Heatmaps from Tweedieverse analyses of extracellular matrix molecules and interactors (c) and modifiers (d) with an increased effect size in keratinocytes, macrophages, fibroblasts, and adipocytes at 1dpw.

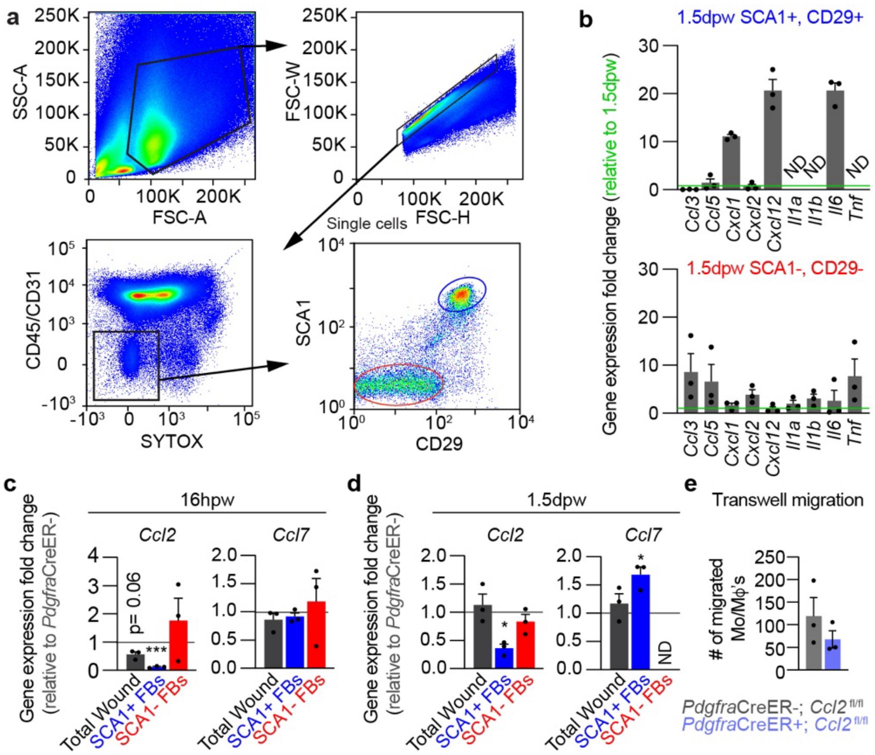


**Supplementary Figure S3. Fibroblast populations exhibit distinct cytokine expression profiles following injury.**

(a) Gating strategy for flow cytometry analysis of mesenchymal populations in skin.

(b) Comparison of cytokine gene expression in the SCA1+, CD29+ and SCA1-, CD29- populations isolated 1.5dpw. Gene expression was measured with qPCR and is normalized to the expression level in 1.5dpw wound beds. *n* = 3 mice per group.

(c-d) Gene expression analysis at 16 hours post-wound (hpw) (c) and 1.5dpw (d) for total wound, SCA1+, or SCA1- fibroblasts from *PdgfraCre*ER+; and *PdgfraCre*ER-; *Ccl2*$^{fl/fl}$ mice. Each bar shows gene expression in the Cre+ group relative to the Cre- group. *n* = 3 mice per group.

(e) Quantification of monocyte (Mo)/macrophage (Mϕ) (CD45+, CD11b+, LY6G-) migration through a transwell towards wound-matched SCA1+ fibroblasts from *Pdgfra*CreER+; or *Pdgfra*CreER-; *Ccl2*$^{fl/fl}$ mice in the bottom of the well. Fibroblasts and monocytes/macrophages were isolated from the same wounds 1.5dpw. *n* = 3 mice per group.

Error bars indicate mean ± SEM. *, $p < 0.05$; ***, $p < 0.001$.

**Supplementary Table S1: List of antibodies**

| Antibody | SOURCE | IDENTIFIER |
|---|---|---|
| APC/Cy7 anti-mouse CD45 rat monoclonal (Clone 30-F11) | Biolegend | Cat# 103116 RRID: AB_312981 |
| FITC anti-mouse CD45 Antibody (Clone 30-F11) | Biolegend | Cat# 103107 RRID: AB_312973 |
| Alexa Fluor 700 anti-mouse CD11b rat monoclonal (Clone M1/70) | Biolegend | Cat# 101222 RRID: AB_493705 |
| eFluor 450 anti-mouse F4/80 rat monoclonal (Clone BM8) | eBioscience | Cat# 48-4801-82 RRID: AB_1548747 |
| BV785 anti-mouse Ly6G rat monoclonal (Clone 1A8) | Biolegend | Cat# 127645 RRID: AB_2566317 |
| PE/Cyanine7 anti-mouse Ly-6G Antibody (Clone 1A8) | Biolegend | Cat# 127618 RRID: AB_1877261 |
| APC-Fire750 anti-mouse CD31 rat monoclonal (Clone 390) | Biolegend | Cat# 102434 RRID: AB_2629683 |
| APC/Fire™ 750 anti-mouse CD326 (Ep-CAM) Antibody (Clone G8.8) | Biolegend | Cat# 118229 RRID: AB_2629758 |
| APC anti-mouse CD326 (Ep-CAM) Antibody (Clone G8.8) | Biolegend | Cat# 118214 RRID: AB_1134102 |
| BV650 anti-mouse Ly-6A/E (SCA1) rat monoclonal (Clone D7) | Biolegend | Cat# 108143 RRID: AB_2629684 |
| Alexa Fluor 700 anti-mouse CD29 Armenian hamster monoclonal (Clone HMbeta1-1) | Biolegend | Cat# 102218 RRID: AB_493711 |
| Anti-CD31 rat monoclonal (Clone MEC13.3) | BD Bioscience | Cat# 550274 RRID: AB_393571 |
| Anti-ER-TR7 rat monoclonal | Abcam | Cat# ab51824 RRID: AB_881651 |
| Anti-F4/80 rat monoclonal (Clone CI:A3-1) | Abcam | Cat# ab6640 RRID: AB_1140040 |
| Anti-ITGA6 rat monoclonal (Clone GoH3) | R&D Systems | Cat# MAB13501 RRID: AB_2128311 |
| Anti-phospho-Histone H3 | Abcam | Cat# ab5176 RRID: AB_304763 |

**Supplementary Table S2: qPCR Primers**

| Gene | Primer sequence (5'-3') | |
|---|---|---|
| *Actb* | Forward | ATCAAGATCATTGCTCCTCCTGAG |
| | Reverse | CTGCTTGCTGATCCACATCTG |
| *Ccl2* | Forward | GTGCTGACCCCAAGAAGGAA |
| | Reverse | GTGCTGAAGACCTTAGGGCA |
| *Ccl3* | Forward | CAG CGA GTA CCA GTC CCT TT |
| | Reverse | GCA GTG GTG GAG ACC TTC AT |
| *Ccl5* | Forward | TGCTCCAATCTTGCAGTCGT |
| | Reverse | GCAAGCAATGACAGGGAAGC |
| *Ccl7* | Forward | CCA TCA GAA GTG GGT CGA GG |
| | Reverse | TGC TTC TTG GCT CCT AGG TTG |
| *Cxcl1* | Forward | TGGCTGGGATTCACCTCAAG |
| | Reverse | CCGTTACTTGGGGACACCTT |
| *Cxcl2* | Forward | CACTCTCAAGGGCGGTCAAA |
| | Reverse | TGGTTCTTCCGTTGAGGGAC |
| *Cxcl12* | Forward | AAC ACA AGA TCC GGC AGA GG |
| | Reverse | ACG GCT AGG AAA GGG TCT CT |
| *Il1a* | Forward | TTGGTTAAATGACCTGCAACA |
| | Reverse | GAGCGCTCACGAACAGTTG |
| *Il1b* | Forward | TTGACGGACCCCAAAAGAT |
| | Reverse | GAAGCTGGATGCTCTCATCTG |
| *Il6* | Forward | AGCCCACCAAGAACGATAGTC |
| | Reverse | TTGTGAAGTAGGGAAGGCCG |
| *Il33* | Forward | CACATTGAGCATCCAAGGAA |
| | Reverse | AACAGATTGGTCATTGTATGTACTCAG |
| *Tnf* | Forward | AAGAGGCACTCCCCCAAAAG |
| | Reverse | ATCCCTTTGGGGACCGATCA |
